# The accumulation of N6-methyl-2’-deoxyadenosine in DNA drives activity-induced gene expression and is required for the extinction of conditioned fear

**DOI:** 10.1101/059972

**Authors:** Xiang Li, Qiongyi Zhao, Wei Wei, Quan Lin, Christophe Magnan, Michael R. Emami, Luis E. Wearick da Silva, Thiago W. Viola, Paul R. Marshall, Jaiyu Yin, Sachithrani U. Madugalle, Sara Nainar, Cathrine Broberg Vågbø, Laura J. Leighton, Esmi L. Zajaczkowski, Ke Ke, Rodrigo Grassi-Oliveira, Magnar Bjørås, Pierre F. Baldi, Robert C. Spitale, Timothy W. Bredy

## Abstract

Here we report that the recently discovered mammalian DNA modification N6-methyl-2’-deoxyadenosine (m6dA) is dynamically regulated in primary cortical neurons, and accumulates along promoters and coding sequences within the genome of activated prefrontal cortical neurons of adult C57/Bl6 mice in response to fear extinction learning. The deposition of m6dA is generally associated with increased genome-wide occupancy of the mammalian m6dA methyltransferase, N6amt1, and this correlates with fear extinction learning-induced gene expression. Of particular relevance for fear extinction memory, the accumulation of m6dA is associated with an active chromatin state and the recruitment of transcriptional machinery to the brain-derived neurotrophic factor (Bdnf) P4 promoter, which is required for Bdnf exon IV mRNA expression and for the extinction of conditioned fear. These results expand the scope of DNA modifications in the adult brain and highlight changes in m6dA as a novel neuroepigenetic mechanism associated with activity-induced gene expression and the formation of fear extinction memory.

## Introduction

In recent years, our understanding of neural plasticity, learning, and memory has been advanced by the demonstration that various epigenetic processes are involved in the regulation of experience-dependent gene expression in the adult brain^1^, and are critically involved in various forms of learning as well as the formation of fear extinction memory^2,3^. DNA methylation, once considered static and restricted to directing cellular lineage specificity during early development, is now recognized as being highly dynamic and reversible across the lifespan^4–6^. Although more than 20 DNA modifications have been identified^7^, nearly all research aimed at elucidating the role of these chemical modifications in the brain has focused on either 5-methylcytosine (5mC) or the more recently rediscovered 5-hydroxymethycytosine (5hmC), which is a functionally distinct oxidative derivative of 5mC^8–10^. 5mC and 5hmC are highly prevalent in neurons relative to other cell types^8,11^. Importantly, both modifications are regulated in response to learning under a variety of conditions, including the extinction of conditioned fear^2,12,13^.

Beyond 5mC and 5hmC, N6-methyl-2’-deoxyadenosine (m6dA) is the most abundant DNA modification in prokaryotes where, in bacteria, m6dA regulates replication, transcription, and transposition^14^. Until recently, m6dA had only been detected in unicellular eukaryotes^15,16^; however, due to significant improvements in mass spectrometry and sequencing technologies, it has now been shown to accumulate in a variety of eukaryotic genomes. For example, in *Chlamydomonas reindardtii*, m6dA is deposited at the transcription start site (TSS) and is associated with increased gene expression^17^ and, in *Drosophila*, the level of m6dA increases across development and is enriched within transposable elements^18^. Furthermore, m6dA appears to be involved in reproductive viability in *Caenorhdoditis elegans*^19^. These observations have led to speculation that m6dA may play an important role in the regulation of gene expression across the lifespan. Several recent studies have extended these findings with the demonstration that m6dA is not only present in the mammalian genome^20–23^, but is also highly dynamic and negatively correlates with LINE retrotransposon activity in both embryonic stem cells and in the adult brain of C57/Bl6 mice following exposure to chronic stress^20,21^. m6dA has also recently been shown to be enriched in gene bodies where it is positively associated with gene expression in human cell lines^22^. However, little is known about the functional relevance of m6dA in specific neuronal populations in the mammalian brain, and a role for m6dA in the gene expression underlying learning and memory has yet to be reported.

The inhibition of learned fear is an evolutionarily conserved behavioral adaptation that is essential for survival. This process, known as fear extinction, involves the rapid reversal of the memory of previously learned contingencies, and depends on gene expression in various brain regions, including the infralimbic prefrontal cortex (ILPFC). The paradigm of fear extinction has long been recognized as an invaluable tool for investigating the neural mechanisms of emotional learning and memory, and the important contribution of the ILPFC to extinction has been demonstrated^23^. A variety of epigenetic mechanisms in the ILPFC have been implicated in the extinction of conditioned fear^2,3,24^ and this behavioral model provides a robust means to interrogate the role of epigenetic mechanisms in a critically important memory process. We therefore set out to explore the role of m6dA within the ILPFC and to elucidate whether it is involved in the formation of fear extinction memory.

## Results

### m6dA is dynamically regulated in response to neuronal activation

Given that the enrichment of chemical modifications in neuronal DNA, including 5mC and 5hmC, confers control over gene expression programs related to cellular identity during early development and in differentiated neurons, and that in lower eukaryotes and humans m6dA has recently been shown to be positively associated with transcription^17,22^, we hypothesized that m6dA may also be fundamental for governing activity-induced gene expression in differentiated neurons and in the adult brain. Three orthogonal approaches were used to establish the presence and dynamic nature of m6dA in cortical neurons (Fig 1A). First, we provided evidence in favor of m6dA as a modified base in neuronal DNA using a gel shift assay on genomic DNA derived from primary cortical neurons that had been treated with DpnI, a bacterially-derived restriction enzyme which cuts double-stranded DNA specifically at methylated adenines predominantly within the GATC motif^17,25^ (Fig. 1B). We then performed liquid chromatography-tandem mass spectrometry (LC-MS/MS) to quantify the global level of m6dA within cortical neurons in response to neural activity. A standard KCl-induced depolarization protocol was used to induce neuronal activity *in vitro*^26^, revealing a significant accumulation of m6dA (Fig. 1C). Finally, an immunoblot using an antibody that recognizes m6dA was used to verify the presence of m6dA following neuronal activation, again revealing a significant increase in m6dA (Fig. 1D). Together, these data demonstrate that m6dA is both a prevalent and inducible DNA modification in primary cortical neurons, findings which are in agreement with recent studies showing that m6dA is an abundant DNA modification in the mammalian genome^22^, is responsive to stress^21^, and is dynamically regulated in human disease states^22^.

**Fig. 1.**
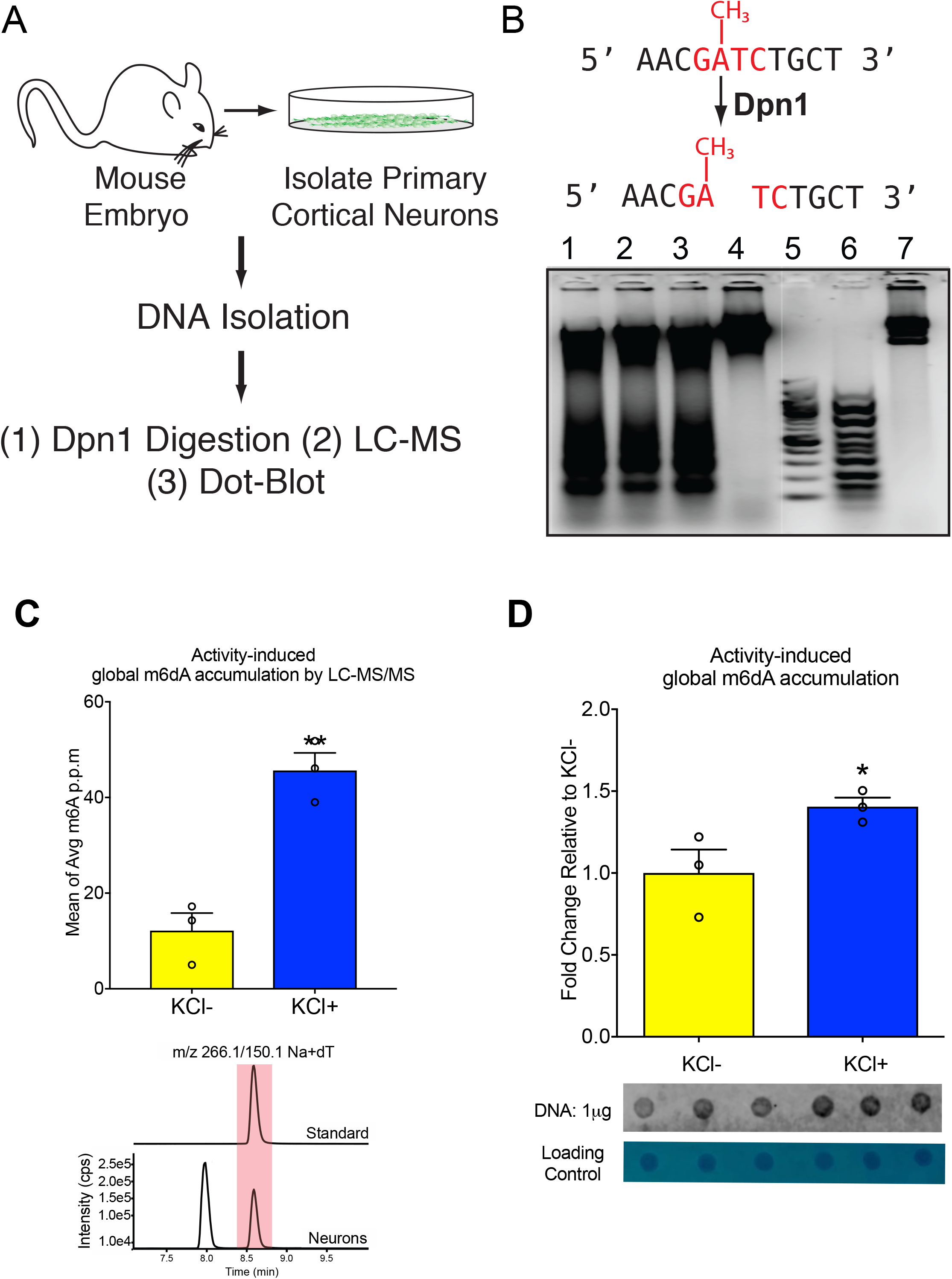
m6dA is present in the neuronal genome and accumulates in response to neural activation. (A) Experimental plan to determine whether m6dA is a functionally relevant base modification in neurons. (B) The Dpn1 enzyme cuts DNA specifically at methylated adenine in GATC linker sequences; Dpn1 digestion reveals the abundance of m6dA in DNA derived from primary cortical neurons, but not in DNA from liver (from the left; lanes 1-3: Dpn1 digested DNA from mouse primary cortical neurons, lane 4: Dpn1 digested DNA from mouse liver, lane 5: DNA ladder, lane 6: Dpn1 digested DNA from *E. coli*, lane 7: undigested DNA from *E. coli*. (C) LC-MS/MS detects a neuronal activity-induced global m6dA induction (7DIV, 20mM KCl, 7 h, two-tailed, unpaired student’s t test, t=6.411, df=4, **p<.01); Representative LC-MS/MS chromatograms: control compound (m6dA standard) and isolated RNase-treated gDNA samples, which were extracted from primary cortical neurons, were used to directly quantify the global level of m6dA. (D) Dot blot assay shows global accumulation of m6dA in stimulated primary cortical neurons (7DIV, 20mM KCl, 7 h, two-tailed, unpaired student’s t test, t=2.634, df=4, *p<.05). (All n=3/group; Error bars represent S.E.M.)

### m6dA accumulates in an experience-dependent manner in the adult brain

Using activity-regulated cytoskeleton associated protein (Arc) and a neuronal nuclear marker (NeuN) as tags for whole-cell fluorescence-activated cell sorting (FACS), we enriched for a specific population of neurons in the mouse ILPFC that had been selectively activated by extinction learning (Suppl. Fig.1A-C). The DpnI-seq approach^27^ was then used to map the extinction learning-induced genome-wide accumulation of m6dA at single base resolution, *in vivo*. As expected, we found that the majority of m6dA sites cleaved by DpnI contained the motif GATC (Suppl. Fig. 2A-B), which is in line with previous reports^28,29^. Specifically, 2,033,704 G(m6dA)TC sites common to both extinction-trained and retention control groups were detected, with 306,207 sites being unique to the extinction training group and 212,326 sites unique to the retention control group (Fig. 2A). Overall, this represents 0.16% of total adenines and, remarkably, 30.49% of all GATCs in the mouse genome, which is significantly more than recent estimates of m6dA in human DNA derived from cell lines^22^ where the predominant motif was [G/CAGG[C/T], and almost an order of magnitude larger than the estimate of differential m6dA regions in DNA derived from the mouse PFC following exposure to chronic stress as assessed by m6dA-immunoprecipitation^21^. Alternating GATC sequences are abundant in eukaryotic DNA and is estimated to account for approximately 0.5% of the total mammalian genome^30,31^. The GATC motif has also been shown to be frequently located at promoter regions where it is directly implicated in gene regulation^32–34^. Our data add to these observations and are the first to demonstrate that m6dA accumulates in neurons that have been selectively activated by fear extinction learning, further suggesting that the dynamic accumulation of G(m6dA)TC may serve a critically important functional role in the epigenetic regulation of experience-dependent gene expression in the adult brain.

**Fig. 2.**
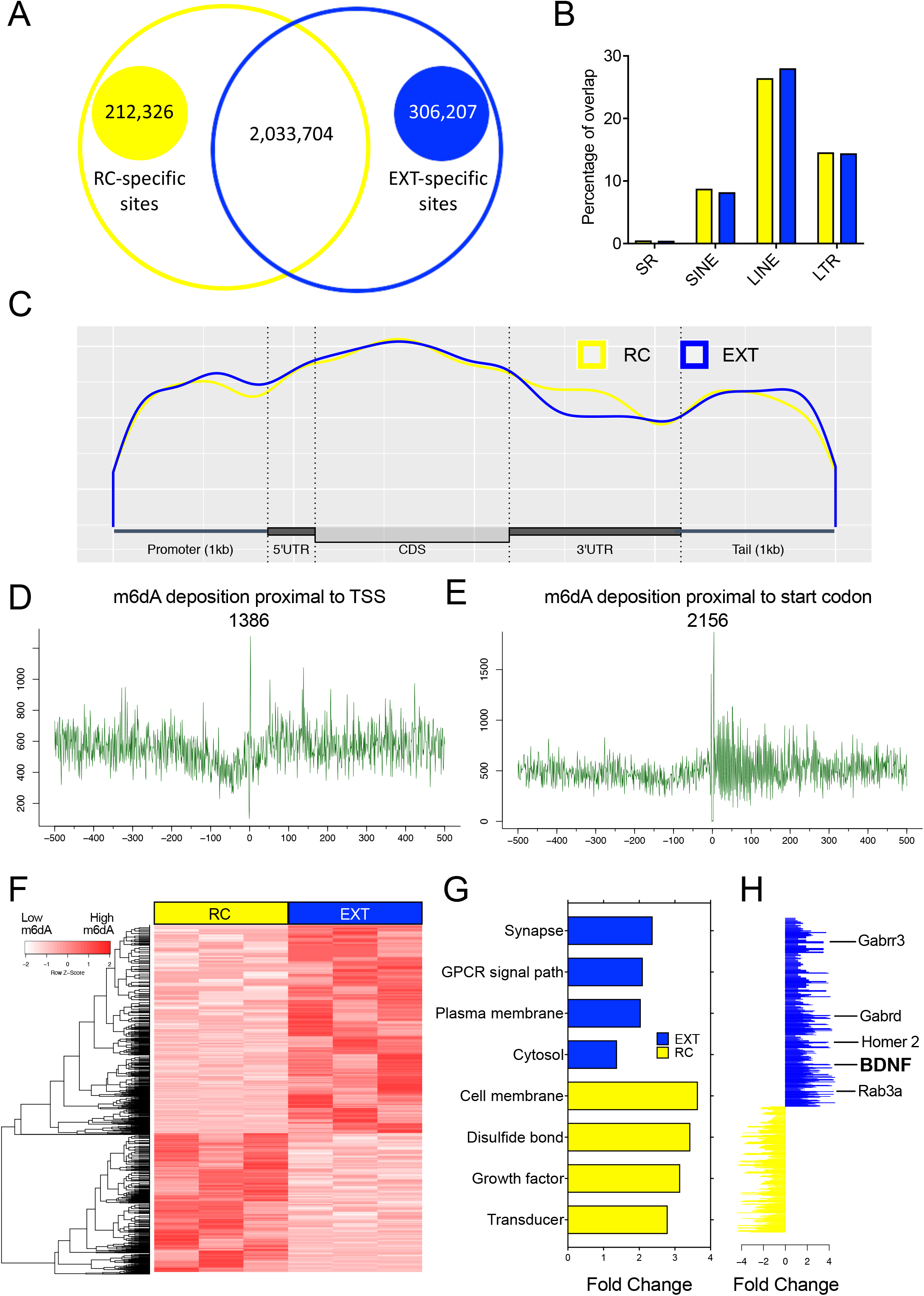
Experience-dependent redistribution of m6dA deposition within ILPFC neurons that have been activated by extinction learning. (A) The Venn diagram shows the learning-induced increase in m6dA sites in retention control (RC) and extinction-trained (EXT) mice. (B) There is no significant difference between RC and EXT groups with respect to the accumulation of m6dA within repetitive elements. (C) Metagene plot shows that m6dA deposition is primarily located in the promoter, 5’UTR and CDS regions. (D) Frequency plots demonstrating that m6dA is enriched at +1bp from TSS, and (E) exhibits a significant increase in deposition at +4bp from the start codon. (F) Representative heat map of genome-wide m6dA enrichment within active neurons after behavioral training. (G) Gene ontology analysis of 10 gene clusters associated with m6dA deposition in RC vs. EXT groups. (H) Representative list of genes that exhibit a significant increase in the accumulation of m6dA and that have been associated with synaptic function, learning and memory.

It has been shown that chronic stress leads to the accumulation of m6dA within LINE1 elements in the adult PFC^23^. Based on this observation, we next examined whether the presence of extinction learning-induced m6dA in activated neurons also overlaps with repeat elements across the genome; however, we found no relationship between the two (Fig. 2B). On the contrary, there was a significant effect of fear extinction learning on the accumulation of m6dA within the promoter, 5’untranslated region (UTR) and exonic coding sequences (CDS) (Fig. 2C). These findings are in accordance with previous studies identifying gene promoters and the trranscription start site (TSS) as critical loci for the dynamic accumulation of m6dA^17,18,27^, as well as the recent discovery of m6dA within coding regions of mammalian DNA^22^. A closer examination of the pattern of m6dA revealed a highly significant increase in the accumulation of m6dA at a site +1bp downstream of the TSS (Fig. 2D), and a sharp increase in m6dA deposition +4bp from the start codon (Fig. 2E). We also detected a significant difference in the experience-dependent accumulation of m6dA between extinction-trained mice and retention controls (Fig. 2F). From a total of 2839 differentially methylated m6dA sites, 1774 GATC sites were specific to extinction. A gene ontology analysis revealed that the most significant cluster specific to the extinction group was for “synapse” (Fig. 2G), with the top synapse-related genes that exhibited a significant accumulation of m6dA in response to extinction learning having previously been shown to be involved in learning and memory (Fig. 2H). Several of these candidates, including Bdnf, Homer2, Gabrr3, Gabrd and Rab3a, were subsequently confirmed in an independent biological cohort using DpnI treatment followed by quantitative PCR, which is represented by a reduced PCR signal when there is more m6dA at a given locus. We again found that a fear extinction learning-induced accumulation of m6dA occured at these genes, but only within neurons that had been activated by extinction training and not within quiescent neurons derived from the same brain region and from the same animals (Suppl. Fig. 3A-H).

To further investigate the relationship between the dynamic accumulation of m6dA and cell type-specific gene expression, RNA-seq was performed on RNA derived from activated and quiescent neurons immediately following fear extinction training. As expected, there was a general increase in gene expression within activated neurons but not in quiescent neurons derived from the same brain region (Fig. 3A). A gene ontology analysis of extinction learning-induced genes again revealed significant extinction learning-related gene clusters, including “synapse”, “dendrite”, and “postsynaptic membrane” (Fig. 3B) with a positive correlation being observed between the accumulation of m6dA and gene expression within neurons selectively activated by fear extinction learning (Fig. 3C). Together with the fact that m6dA was only apparent in active neurons, our data indicate that the dynamic accumulation of m6dA governs different aspects of gene regulation and, in the case fear extinction learning, it is permissive and occurs in a cell-type- and state-dependent manner.

**Fig. 3.**
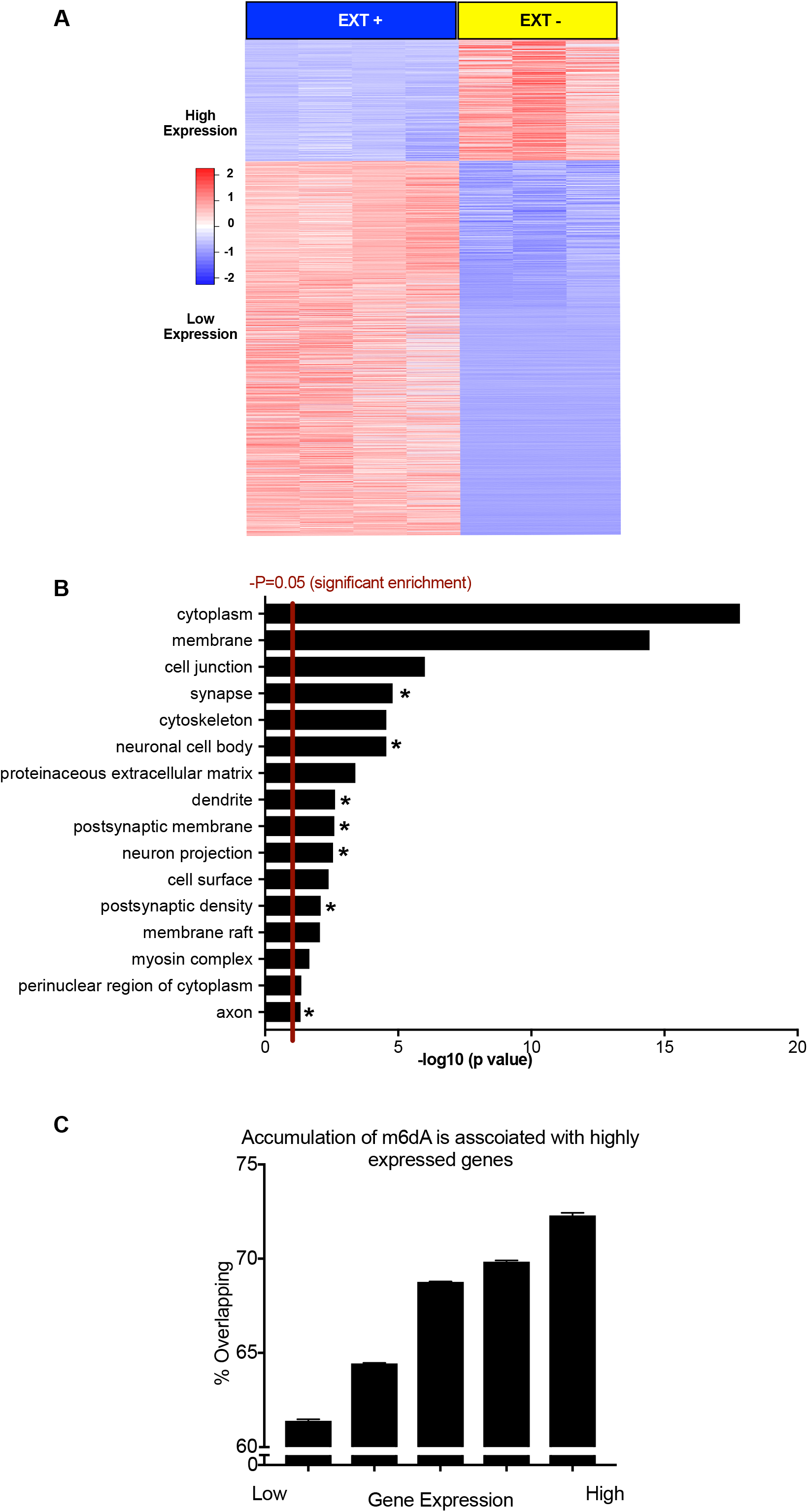
Extinction learning-induced accumulation of m6dA positively correlates with gene expression in activated neurons. (A) Representative heat map of mRNA expression within quiescent neurons (EXT-) vs activated neurons (EXT+). (B) Gene ontology analysis shows gene clusters enriched in the upregulated and differentially expressed genes; neuronal activity-related gene clusters are highlighted. (C) Extinction learning-induced m6dA sites positively correlate with highly expressed genes.

### The expression of N6amt1 is activity-dependent and its deposition is associated with extinction learning-induced changes in m6dA

N6-adenine-specific DNA methyltransferase 1 (N6amt1) was originally described as a mammalian ortholog of the yeast adenine methyltransferase MTQ2. Homologs of N6amt1 have been shown to methylate N6-adenine in bacterial DNA^35^, and mammalian N6amt1 has been shown to be a glutamine-specific protein methyltransferase^36^. N6amt1 is expressed in the mouse neocortex (http://mouse.brain-map.org/experiment/show?id=1234), as is N6amt2, which shares a highly conserved methyltransferase domain (http://mouse.brain-map.org/experiment/show?id=69837159). In order to obtain deeper insight into the underlying mechanism by which m6dA accumulates in the mammalian genome and regulates gene expression, we first examined the expression of N6amt1 and N6amt2 in primary cortical neurons *in vitro* and in the adult ILPFC in response to fear extinction learning. N6amt1 exhibited a significant increase in mRNA expression in primary cortical neurons in response to KCl-induced depolarization (Suppl.Fig. 4A), whereas there was no effect on N6amt2 (Suppl.Fig. 4B). We next sought to determine whether the effects observed in primary cortical neurons also occur in the adult brain by examining N6amt1 and N6amt2 mRNA expression in the ILPFC in extinction-trained mice relative to retention controls. Similar to the effect of KCl-induced depolarization on m6dA accumulation and N6amt1 gene expression *in vitro*, fear extinction training led to a significant increase in N6amt1 mRNA expression in the ILPFC (Fig. 4A), again with no detectable change in N6amt2 (Fig. 4B). There was also a concomitant increase in N6amt1 protein expression in the ILPFC (Fig. 4C) with no effect on the level of N6amt2 (Fig. 4D), suggesting that N6amt1 is selectively induced and is a potentially important epigenetic modifier that mediates the accumulation of m6dA in the adult brain in response to fear extinction learning.

**Fig. 4.**
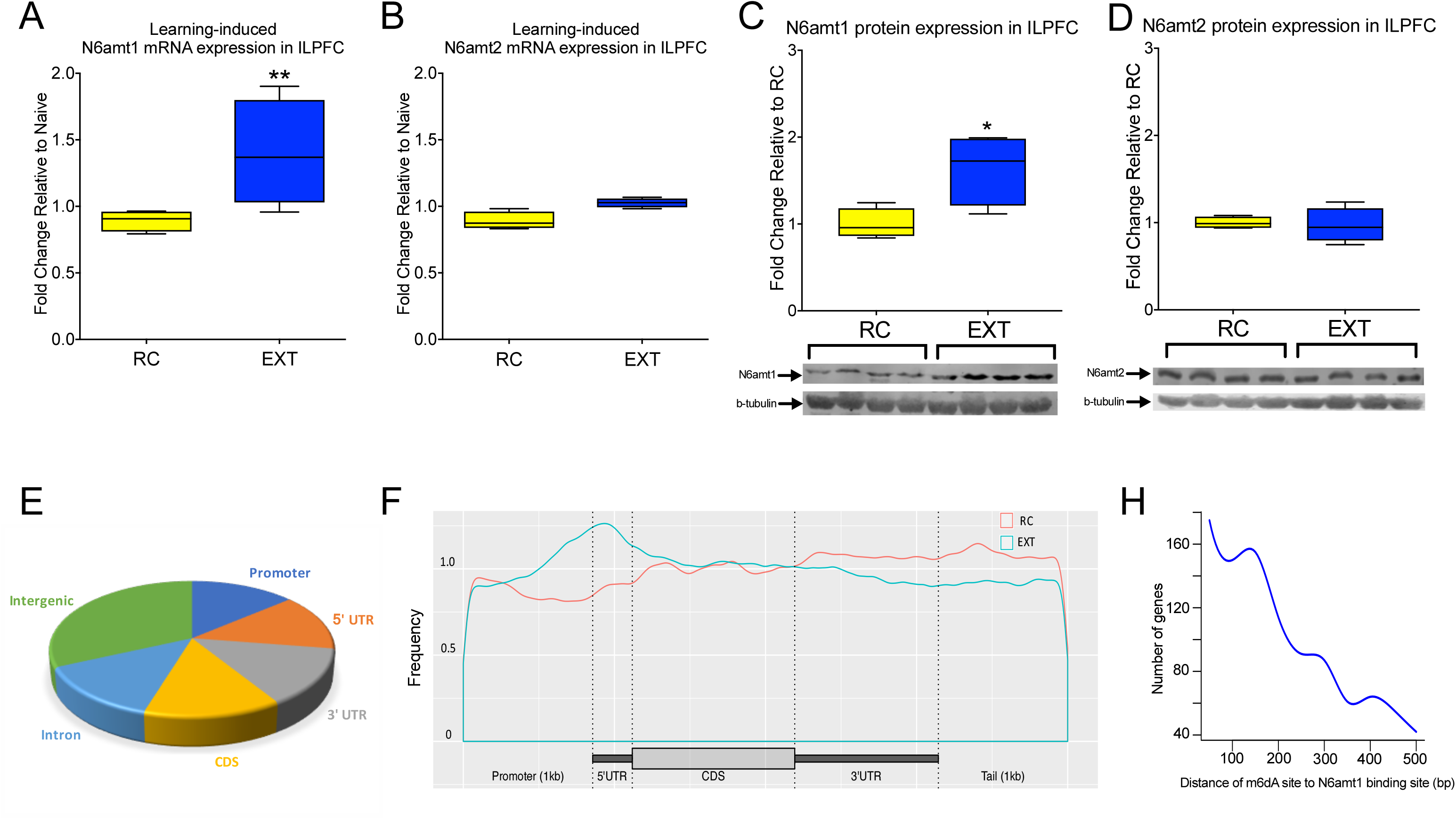
N6amt1 mRNA expression is induced in the ILPFC in response to fear extinction learning, and N6amt1 occupancy increases within gene promoters and 5’UTR. (A) Extinction-learning leads to increased expression of N6amt1 in the ILPFC (two-tailed, unpaired student’s t test, t=2.483, df=6, **p<.01). (B) No significant effect of learning on N6amt2 mRNA expression in the ILPFC. (C-D) N6amt1 protein level is induced post to extinction but not N6amt2 (two-tailed, unpaired student’s t test, t=2.843, df=6, *p<.05). (E) N6amt1 distribution across genome. (F) Extinction training-induced increased in N6amt1 occupancy at promoter and 5’UTR regions. (H) There is a positive relationship between N6amt1 deposition and m6dA sites within promoter and 5’UTR region with more genes showing N6amt1 within 0-200bp of m6dA sites.

To extend our understanding of the role of N6amt1 in fear extinction, we next performed N6amt1 chromatin immunoprecipitation sequencing (ChIP-seq) on samples derived from the ILPFC of fear extinction-trained mice. We found that although N6amt1 occupancy was equally distributed across the genome (Fig. 4E), there was a significant increase in N6amt1 occupancy around gene promoters and 5’UTR in response to fear extinction learning (Fig. 4F). Intriguingly, we found 995 genes that exhibit an increase both N6amt1 depostion and an accumulation of m6dA surrounding the TSS. This represents over 72% of the total number of highly expressed genes in the fear extinction group that show a correlated increase in m6dA (Suppl.Table.1). In addition, a distribution plot revealed a strong association between N6amt1 binding sites and m6dA sites (Fig. 4H). These findings imply a functional relationship between an extinction learning-induced increase in N6amt1 occupancy and the accumulation of m6dA in the adult brain. However, the incomplete nature of the overlap between the two datasets also suggests that there are yet to be identified epigenetic modifiers which contribute to the dynamic accumulation of m6dA, and that these may be associated with other factors such as the temporal dynamics of N6amt1 recruitment following learning.

In an effort to establish a functional relationship between N6amt1 and the accumulation of m6dA in neuronal DNA, we next performed an N6amt1 overexpression experiment on primary cortical neurons, *in vitro*. Compared with a scrambled control, there was a global increase in m6dA within the cells that overexpressed the full length N6amt1 (Suppl. Fig. 5A-B). Moreover, by knockdown N6amt1 *in vitro*, we observed an reduction of m6dA (Suppl. Fig. 5C-D). These findings are in agreement with the recent confirmation of catalytically active N6amt1 in mammalian DNA, which was shown be necessary for the conversion of adenine to m6dA under both synthetic conditions and in human cell lines^22^. Importantly, in that study, there was no effect of N6amt1 on the acccumulation of RNA m6A, therefore verifying a functional role for N6amt1 as a mammalian DNA adenine methyltransferase.

### Extinction learning-induced N6amt1-mediated accumulation of m6dA drives Bdnf exon IV mRNA expression in the ILPFC

Bdnf is the most widely expressed inducible neurotrophic factor in the central nervous system^37^, and is directly involved in extinction-related learning and memory^38^. In the adult brain, the accumulation of 5mC within Bdnf gene promoters is altered by experience^39^, and this epigenetic mechanism appears to be necessary for the regulation of gene expression underlying remote memory^12^. The Bdnf locus comprises at least eight homologous noncoding exons that contribute to alternate 5’-UTRs, and a ninth that contributes a protein coding sequence and 3’-UTR^40,41^. The complex structure of this genomic locus has led to the idea that Bdnf mRNA expression may be driven by DNA modifications that guide distinct sets of transcription factor complexes to initiate the transcription of the various isoforms^42^, all of which could be important for learning and memory. This is supported by the fact that exon IV is highly activity-dependent and plays a direct role in the formation of fear extinction memory^24,43^.

We observed a highly specific accumulation of m6dA at a GATC site immediately downstream of the TSS of the Bdnf P4 promoter in fear extinction-trained mice (Suppl Fig 3A and Fig. 5A). DNA immunoprecipitation analysis using an m6dA-specific antibody confirmed that extinction training led to an increase in m6dA at this locus and that this signal could be detected within a mixed homogenate derived from the ILPFC of extinction-trained mice (Fig. 5B). This is the only GATC site found within 500bp of the Bdnf exon IV TSS, which again suggests a high level of selectivity with respect to where and when m6dA dynamically accumulates in the genome in response to extinction training. Formaldehyde-assisted isolation of regulatory elements (FAIRE) followed by qPCR was then used to determine chromatin status ^44, 45^ and increased activity at this m6dA-modified GATC site was revealed (Fig 5C). When considered in conjunction with the increased deposition of H3K4^me3^ (Fig. 5D), this further suggests a functionally relevant relationship between m6dA and the induction of an open chromatin state. A consensus sequence for the activating transcription factor Yin-Yang (YY1)^23,46,47^ is located adjacent to the m6dA site, and fear extinction learning led to a significant increase in the recruitment of YY1 (Fig 5E), as well as components of the transcription machinery, including TFIIB (Fig. 5F) and Pol II (Fig 5G). The activity-induced changes in N6amt1 occupancy, m6dA accumulation and related effects on the local chromatin landscape and transcriptional regulators strongly correlated with increased Bdnf exon IV mRNA expression in response to fear extinction training (Fig. 5H). There was no effect in IgG controls (Suppl. Fig. 6A-F), at a distal GATC motif located 1000bp upstream of the TSS (Suppl. Fig. 7A-E), or proximal to the TSS in the Bdnf P1 promoter (Suppl. Fig. 8A-H). The proximal promoter region of another plasticity-related gene, Rab3A, also exhibited a pattern of epigenetic modification similar to that of Bdnf P4, and a subsequent increase in gene expression in response to fear extinction learning (Supp Fig 9A-F). This further suggests that m6dA has a generalized role in the epigenetic regulation of experience-dependent gene expression in the adult brain.

**Fig. 5.**
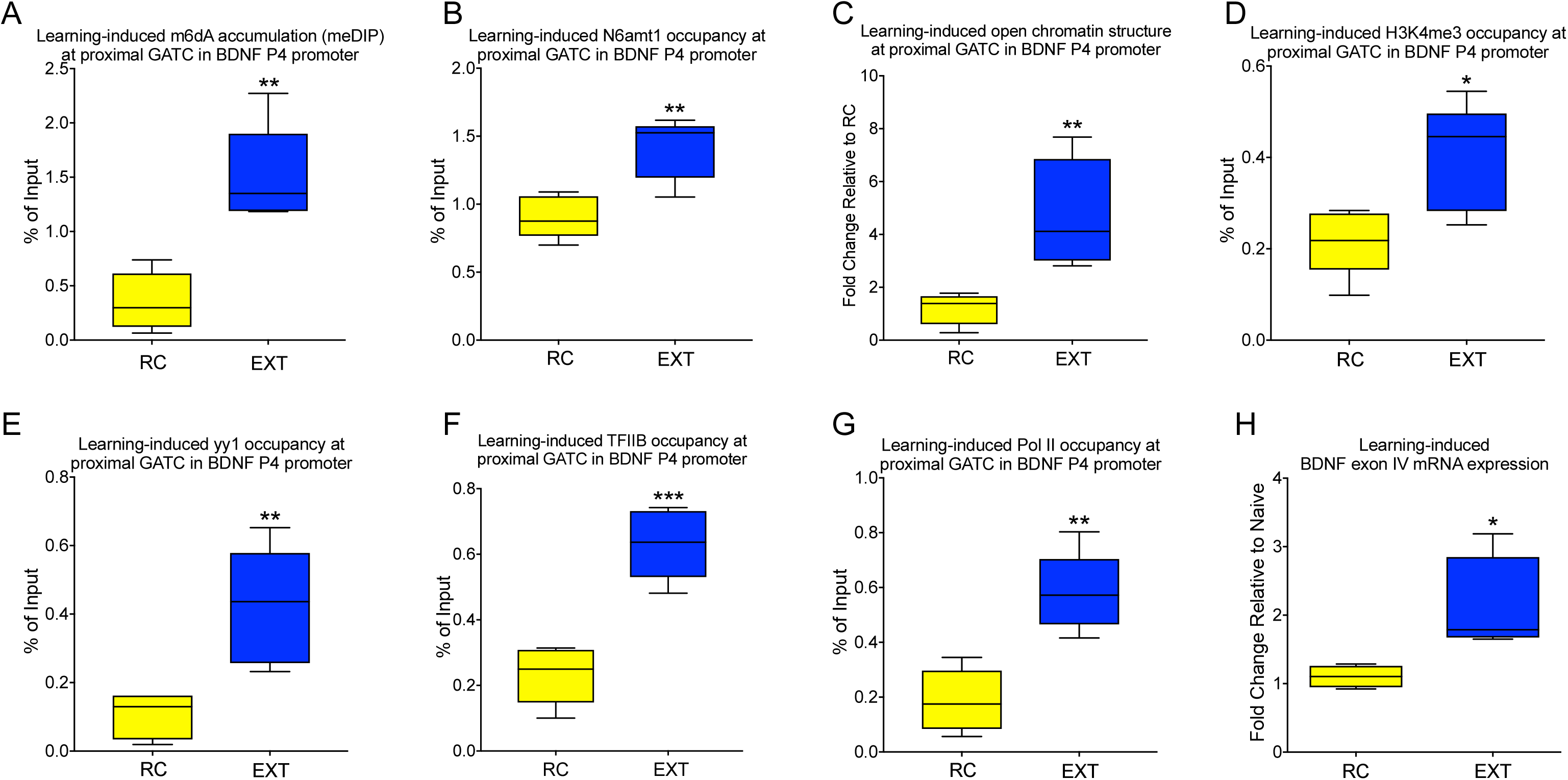
Extinction learning-induced accumulation of m6dA is associated with an active chromatin landscape and increased bdnf exon IV mRNA expression. Fear extinction learning (EXT), relative to fear conditioned mice exposed to a novel context (retention control: RC), led to (A) increased m6dA at the previously identified GATC site (two-tailed, unpaired student’s t test, t=4.921, df=8, **p<.01), (B) a selective increase in N6amt1 occupancy (two-tailed, unpaired student’s t test, t=4.133, df=8, **p<.01), (C) a increased open chromatin structure was detected by using FAIRE-qPCR (two-tailed, unpaired student’s t test, t=3.76, df=8, **p<.01), (D) a significant increase in H3K4^me3^ occupancy (two-tailed, unpaired student’s t test, t=2.986, df=8, *p<.05), (E) an increase in the recruitment of YY1 (two-tailed, unpaired student’s t test, t=3.885, df=8, **p<.01), (F) an increase in TFIIB occupancy (two-tailed, unpaired student’s t test, t=6.474, df=8, **p<.01), (G) an increase in Pol II occupancy (two-tailed, unpaired student’s t test, t=4.838, df=8, **p<.01). (H) a significant increase in bdnf exon IV mRNA expression within the ILPFC (two-tailed, unpaired student’s t test, t=2.685, df=6, *p<.05). (All n=5/group, Error bars represent S.E.M.)

### N6amt1-mediated accumulation of m6dA is associated with increased gene expression and the formation of fear extinction memory

Having established a relationship between the fear extinction learning-induced accumulation of m6dA and the regulation of Bdnf exon IV mRNA expression *in vivo*, we next investigated whether lentiviral-mediated knockdown of N6amt1 in the ILPFC affects the formation of fear extinction memory. We first validated the efficiency of the knockdown construct *in vivo*, revealing excellent transfection efficiency and a reliable decrease in N6amt1 mRNA expression when the construct was infused directly into the ILPFC prior to behavioral training (Fig. 6A-B). There was no effect of N6amt1 shRNA on within-session performance during the first 10 conditioned stimulus exposures during fear extinction training (Fig. 6C-D), and there was no effect of N6amt1 shRNA on fear expression in mice that had been fear conditioned and exposed to a novel context without extinction training (Fig. 6E-left, pre-CS). However, we observed a highly significant impairment in fear extinction memory in mice that had been extinction trained in the presence of N6amt1 shRNA (Fig. 6E-right). Infusion of N6amt1 shRNA into the prelimbic region of the PFC, a brain region immediately dorsal to the ILPFC predominantly involved in fear acquisition, had no effect on extinction memory (Suppl. Fig. 10A-C). These data imply a critical role for the N6amt1-mediated accumulation of m6dA in the ILPFC in the regulation of regulating fear extinction memory formation as opposed to non-specific negative effects on fear-related learning and memory.

**Fig. 6.**
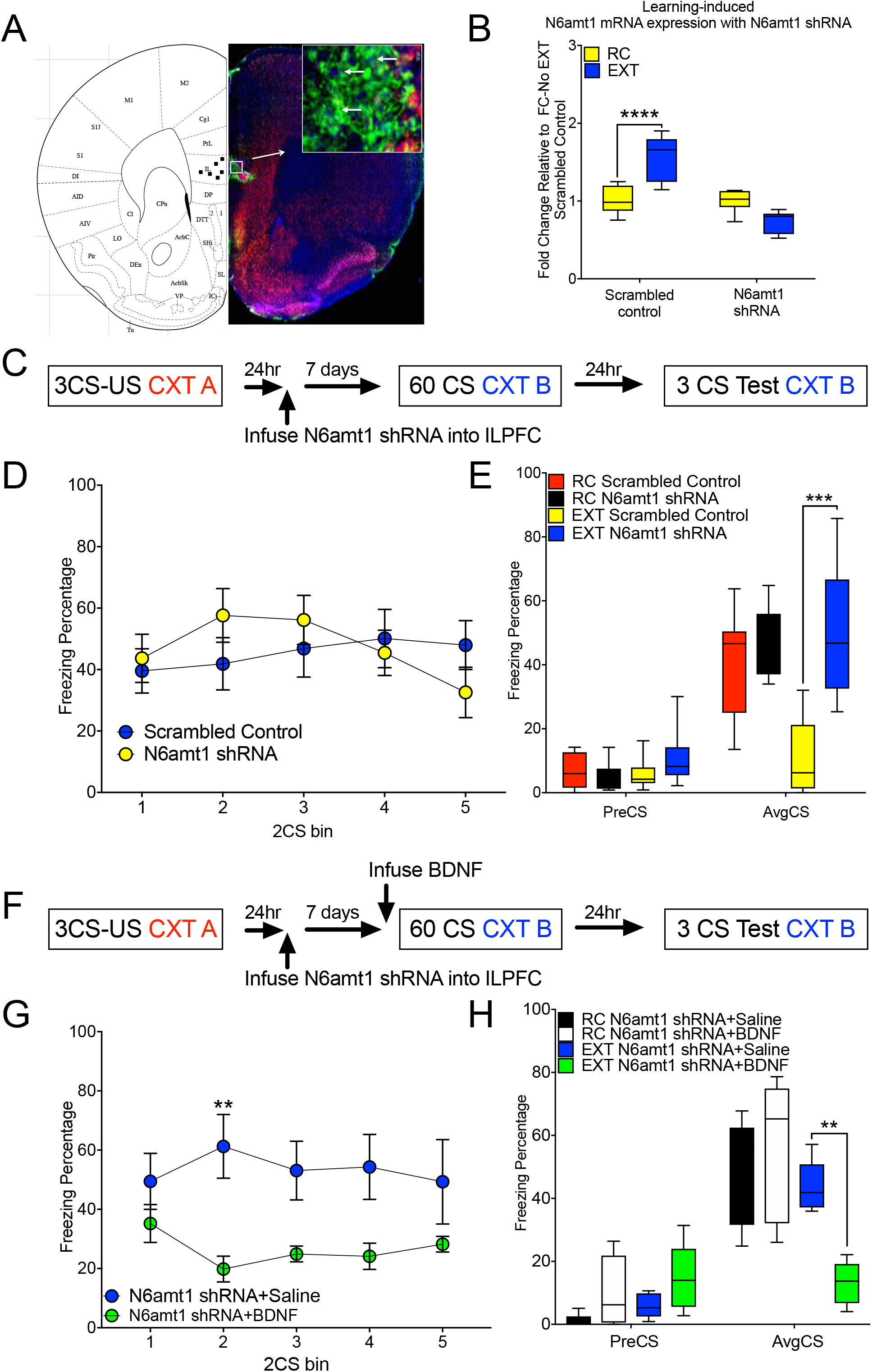
N6amt1-mediated accumulation of m6dA is required for fear extinction memory and for learning-induced bdnf exon IV mRNA expression in the ILPFC. (A) Left: representative image of cannula placement in the ILPFC, Right: transfection of N6amt1 shRNA into the ILPFC. (B) N6amt1 shRNA blocks the induction of N6amt1 mRNA expression following extinction learning (two-way ANOVA, F_1,12_=11.44, p<.01; Tukey’s posthoc test: Scrambled control EXT vs. N6amt1 shRNA EXT, ***p<.001). (C) Schematic of the behavioral protocol used to test the effect of lentiviral-mediated knockdown of N6amt1 in the ILPFC on fear extinction memory. (D) There was no effect of N6amt1 shRNA on within-session performance during the first 10 conditioned stimulus exposures during fear extinction training. (E) Although there was no effect of N6amt1 shRNA on fear expression in mice that had been fear conditioned and exposed to a novel context without extinction training, N6amt1 knockdown led to a significant impairment in fear extinction memory (*p<.05). (F) Schematic of the behavioral protocol used to test the effect of BDNF Injection within N6amt1 knockdown animals on fear extinction memory. (G) ILPFC infusion of BDNF has minimum effect during the section of extinction training (two-way ANOVA, F_1,40_=25.34, p<.0001; Sidak’s posthoc test: N6amt1 shRNA+Saline vs. N6amt1+BDNF, **P<.01) promotes extinction rescues the N6amt1 shRNA-induced impairment in fear extinction memory (two-way ANOVA, F_3,35_=4.749, p<.01; Tukey’s posthoc test: EXT N6amt1 shRNA+Saline vs. EXT N6amt1+BDNF, **p<.01). (All n=6-8 per group; Error bars represent S.E.M.)

In order to draw stronger conclusions about the relationship between m6dA Bdnf mRNA expression and extinction memory, we next asked whether the direct application of recombinant Bdnf into the ILPFC prior to extinction training would rescue the impairment of extinction memory associated with N6amt1 knockdown. In the presence of N6amt1 shRNA, Bdnf-treated mice exhibited a significant reduction in freezing relative to saline-infused mice during extinction training (Fig 6F-H), which implies a causal relationship between N6amt1-mediated accumulation of m6dA, Bdnf exon IV expression, and the formation of fear extinction memory. With respect to the epigenetic landscape and transcriptional machinery surrounding the Bdnf P4 promoter, knockdown of N6amt1 prevented the fear extinction learning-induced increase in N6amt1 occupancy (Fig. 7A) and the accumulation of m6dA (Fig. 7B). N6amt1 knockdown also blocked the fear extinction learning-induced change in chromatin state (Fig 7C), as well as the previously observed increase in H3K4^me3^ (Fig. 7D), YY1 (Fig. 7E), TFIIB (Fig. 7F) and Pol II (Fig. 7G) recruitment to the Bdnf P4 promoter. Finally, N6amt1 knockdown prevented the fear extinction learning-induced increase in Bdnf exon IV mRNA expression (Fig. 7H). Taken together, these findings indicate that, in the ILPFC, the dynamic accumulation of m6dA is required for the epigenetic regulation of experience-dependent Bdnf exon IV expression and is critically involved in the formation of fear extinction memory.

**Fig. 7.**
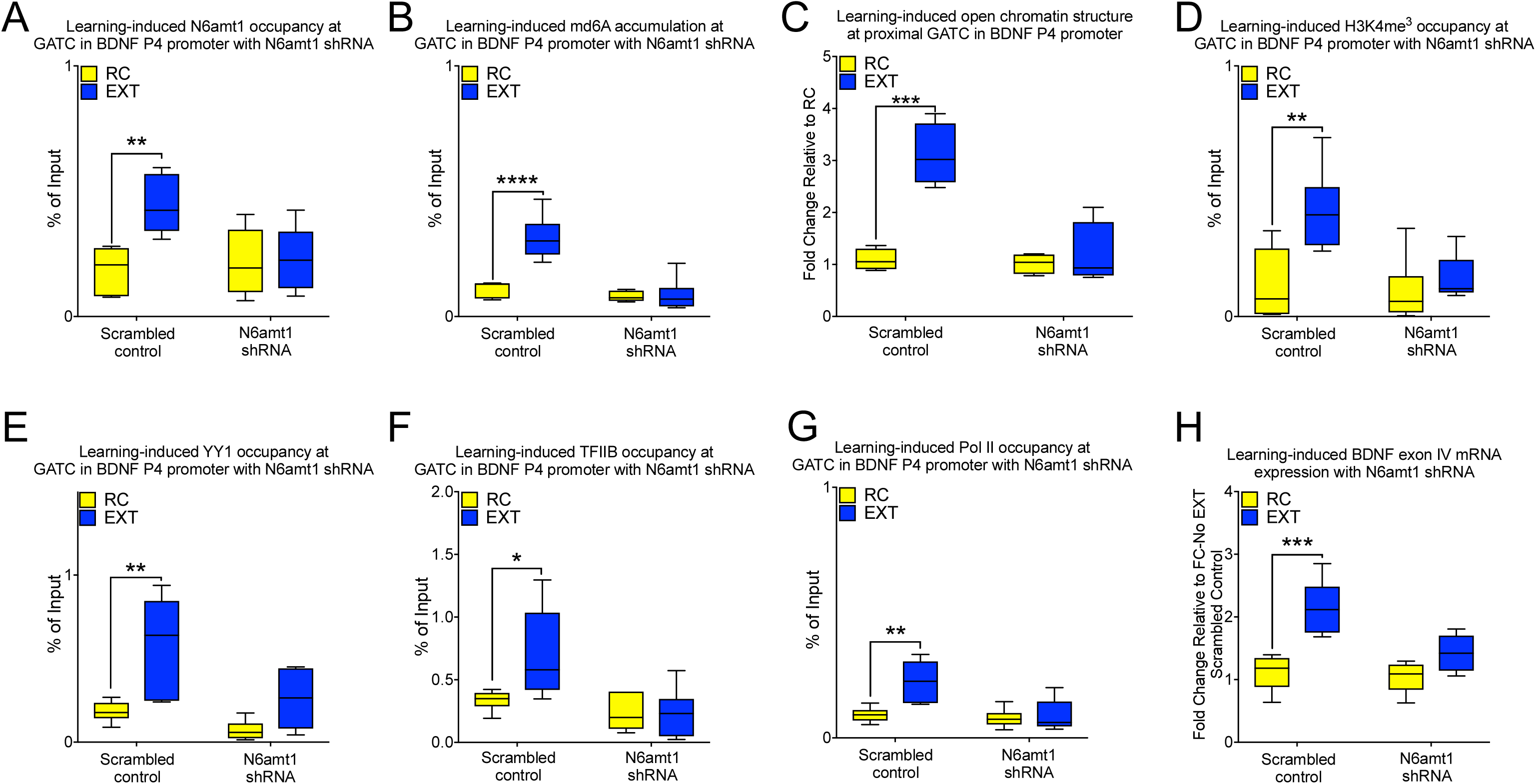
N6amt1 knockdown prevents the learning-induced accumulation of m6dA and related changes in chromatin and transcriptional landscape associated with the BDNF P4 promoter. N6amt1 shRNA blocked (A) the learning-induced increase in N6amt1 occupancy (two-way ANOVA F_1,20_ = 7.663, p<.05; Dunnett’s posthoc test: scrambled control RC vs. scrambled control EXT, **p<.01) and (B) the deposition of m6dA (two-way ANOVA F_1,19_ = 18.56, p<.0001; Dunnett’s posthoc test: scrambled control RC vs. scrambled control EXT, ****p<.0001), or (C) open chromatin structure (two-way ANOVA F_1,12_ = 19.38, p<.001; Dunnett’s posthoc test: scrambled control RC vs. scrambled control EXT, ***p<.0001), (D) the accumulation of H3K4me^3^ (two-way ANOVA F_1,19_ = 9.815, p<.01; Dunnett’s posthoc test: scrambled control RC vs. scrambled control EXT, **p<.01), (E) YY1 (two-way ANOVA F1,20 = 17.64, p<.001; Dunnett’s posthoc test: scrambled control RC vs. scrambled control EXT, **p<.01), (F) induction of TFIIB occupancy (two-way ANOVA F_1,20_ = 10.03, p<.01; Dunnett’s posthoc test: scrambled control RC vs. scrambled control EXT, *p<.05) and (G) RNA Pol II (two-way ANOVA F_1,20_ = 8.883, p<.01; Dunnett’s posthoc test: scrambled control RC vs. scrambled control EXT, **p<.01) occupancy at the proximal GATC site within the BDNF P4 promoter. Also, (H) N6amt1 shRNA inhibited the bdnf exon IV mRNA expression (two-way ANOVA F_1,16_ = 20.95, p<.001; Dunnett’s posthoc test: scrambled control RC vs. scrambled control EXT, ***p<.001). (All n=5-6/group, Error bars represent S.E.M.)

## Discussion

Although more than 20 different base modifications are known to occur in DNA^7^, only 5mC and 5hmC have been studied in any detail within the mammalian brain. Here we provide the first evidence that the learning-induced accumulation of m6dA in post-mitotic neurons is associated with an increase in gene expression and critically involved in the formation of fear extinction memory. m6dA has emerged as a functionally relevant DNA modification that is commonly found in bacterial DNA and lower eukaryotes^15,16,28,29,48,49^. Recently, it has been shown that m6dA is abundant in the mammalian genome^22, 25 21^ and that the accumulation of m6dA in the PFC is associated with chronic stress^21^. Using a combination of HPLC-MS/MS and a dot blot assay, we have extended these observations and provide strong evidence for a global induction of m6dA in response to neuronal activation. In addition, we have revealed an activity-dependent increase in the expression of the m6dA methyltransferase N6amt1, which appears to be critical for the extinction learning-induced accumulation of m6dA at the Bdnf P4 promoter and for the extinction of conditioned fear. Importantly, we have found that the effect of experience on m6dA accumulation occurs only in neurons that have been activated by training and not in quiescent neurons from the same brain region. These data therefore suggest that neurons employ m6dA as an epigenetic regulatory mechanism that is engaged specifically under activity-induced conditions, and that this is mediated by the action of N6amt1. Whether m6dA has similar regulatory control over experience-dependent gene expression in other cell types and in other regions of the brain remains to be determined.

As indicated, full length overexpression of N6amt1 led to a global increase in m6dA within primary cortical neurons, and a positive correlation between N6amt1 occupancy and the level of m6dA was observed, similar to the recent demonstration that catalytically active N6amt1 mediates the accumulation of m6dA in human DNA^22^. However, as indicated by our genome-wide profiling, N6amt1 did not show complete overlap with sites of extinction-learning induced m6dA accumulation within gene promoters (Suppl. Table. 1). Therefore, it is likely that, in order to confer temporally regulated changes in the accumulation of m6dA in cortical neurons, N6amt1 must also work in complex with other factors. This is not an unreasonable assumption, as other methyltransferases such as DNMT1 and Tet2 have been shown to require interaction with scaffolding proteins and/or regulatory RNAs, which then determine their target on DNA^50–52^. Regardless of these underlying mechanisms, there is clearly a functional relationship between N6amt1 and the accumulation of m6dA in response to fear extinction training. Future studies will determine the full repertoire of proteins and RNA that are required to direct N6amt1 to sites of action on DNA in an experience- or activity-dependent manner. We have previously found that the pattern of 5mC within the adult brain differs in neurons and non-neuronal cells^53^ and that 5hmC exhibits a dramatic redistribution in the adult ILPFC in response to fear extinction learning^2^. Together, these lines of evidence suggest that learning-induced changes in DNA modification may be both dynamic and cell-type specific, a conclusion that is supported by our current findings as it is evident that the accumulation of m6dA in post-mitotic neurons relies on their activation state.

It is noteworthy that the fear extinction learning-induced accumulation of m6dA was prominent not only around the TSS, but also along the CDS (Fig 2), reflecting a similar pattern in the human genome^22^. Interestingly, previous work has shown that m6dA is associated with Pol II transcribed genes^ref^ and that the accumulation of m6dA in exons positively correlates with gene transcription^ref^. Together with our findings, these data suggest that m6dA may play an important role in initiating transcription by promoting an active chromatin state and, with the recruitment of Pol II, may contribute to the efficiency of Pol II read-through along the gene body. Moreover, m6dA has been shown to contribute to base flipping^54^, and G(m6dA)TC has been shown to induce structural changes in DNA^55^. During elongation, these effects on DNA structure may maintain the transcription bubble by lowering the energetic barrier for the RNA polymerase active site^56,57^. Interestingly, we found that highly expressed genes tend to have more m6dA within their promoter region (Fig 3E). Furthermore, m6dA has been shown to overlap with nucleosome-free regions^17^, which also serves to facilitate transcription elongation^59,60^. This suggests an essential role for the deposition of m6dA along the CDS in regulating activity/learning-induced transcriptional processes, which are required for the underlying changes in gene expression that accompany the formation of fear extinction memory. Future studies will examine the direct relationship between the dynamic accumulation of m6dA and DNA structure states, and their influence on gene expression and on other forms of learning and memory.

Our data demonstrate that the activity-induced expression of Bdnf exon IV within the ILPFC following behavioral training is functionally related to an N6amt1-mediated increase in the accumulation of m6dA at the Bdnf P4 promoter. This is also associated with an open chromatin structure as well as the presence of H3K4^me3^, an epigenetic mark that reflects an active chromatin state, and accompanied by the increased recruitment of the transcription-related factors YY1 and TFIIB, as well as Pol II (Fig. 5). These findings demonstrate that the accumulation of m6dA surrounding the TSS of the Bdnf P4 promoter drives activity-induced and experience-dependent exon IV mRNA expression, which is in agreement with recent findings on m6dA-mediated transcriptional activation in lower eukaryotes^17^ and in human cell lines^22^.

Other DNA modifications, including oxidative derivatives of 5mC, are found within gene promoters as well as gene bodies, and have been shown to interact with Pol II to induce transient pausing of the Pol II elongation complex to promote gene expression^18,58,59^. This suggests that Pol II has the capacity to detect a variety of DNA modifications, including m6dA, and that this interaction may serve to fine-tune the rate of Pol II-mediated transcription. The high degree of specificity of m6dA accumulation at the specific GATC site proximal to the TSS in the Bdnf P4 promoter further implies tight control over Bdnf exon IV expression through regionally selective epigenetic regulation by this DNA modification. Pol II has been shown to be recruited to the Bdnf P1 promoter in a spatiotemporally regulated manner, resulting in ‘waves’ of Bdnf exon I mRNA expression^60^, which may explain why no change in m6dA was observed at the Bdnf P1 promoter at the timepoint interrogated in this study.

In summary, we have shown that the N6amt1-mediated accumulation of m6dA is dynamically regulated in the mammalian genome and that its deposition drives activity-induced Bdnf exon IV mRNA expression and is required for the extinction of conditioned fear. Our findings suggest a model where, in selectively activated neurons in the adult brain, the accumulation of m6dA serves as a permissive epigenetic signal for the regulation of activity- or learning-induced gene expression (Fig. 8). These results expand the scope of experience-dependent DNA modifications in the brain and stongly indicate that the information-processing capacity of DNA in post-mitotic neurons is far more complex than current perspectives generally appreciate. We predict that a large number of functional modifications on *all four* canonical nucleobases, with diverse roles in the epigenetic regulation of experience-dependent gene expression, learning and memory, remain to be discovered.

**Fig. 8.**
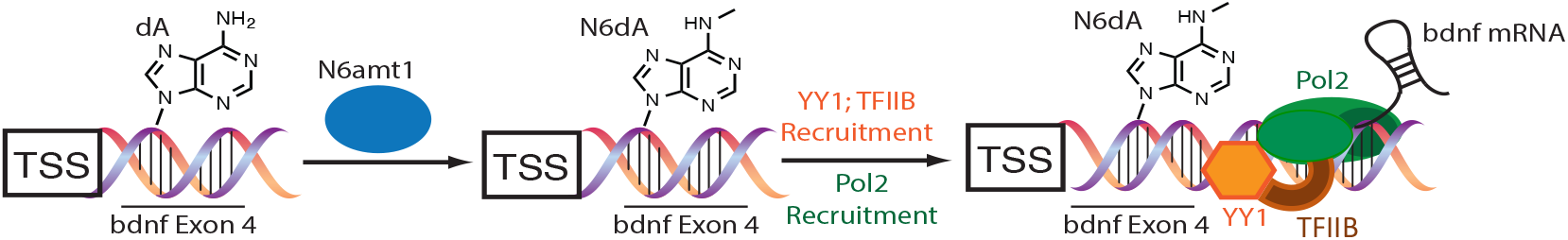
A schematic of the hypothesized role of m6dA within the Bdnf P4 promoter of neurons activated by extinction learning. The dynamic accumulation of m6dA drives activity-dependent Bdnf exon IV gene expression by facilitating an active and open chromatin state, and the recruitment of essential components of the transcriptional machinery, including YY1, TFlIb and PolII.

---

**Suppl. Fig. 1.**
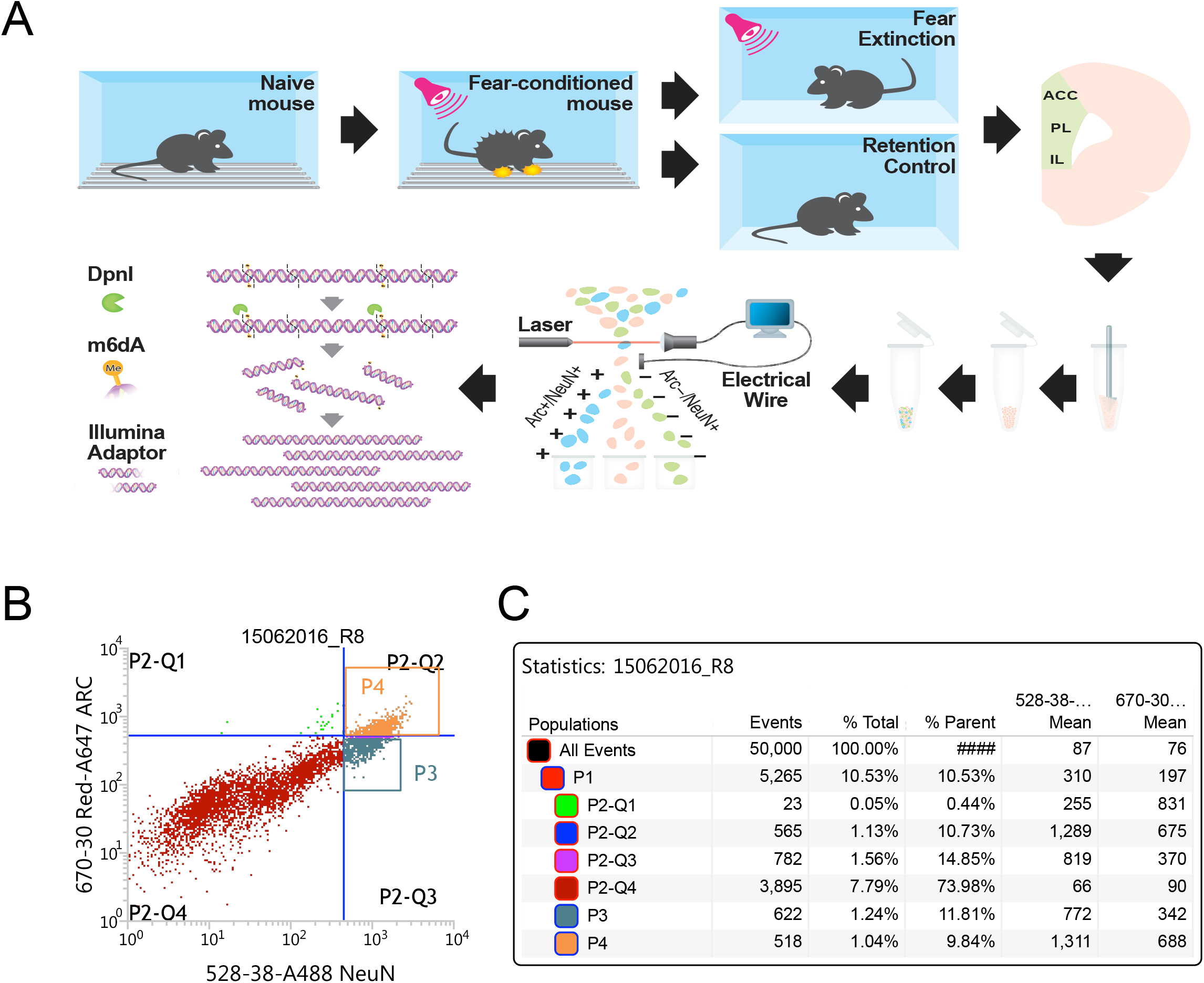
Using flow cytometry coupled with DpnI-seq, we generated a genome-wide profile of cell-type-specific, learning-induced m6dA deposition at single-base resolution. (A) The schema shows the work flow. (B) Flow cytometry scatterplots demonstrating enrichment for activated neurons using Arc and NeuN as tags. (C) FACS reports show there is distinct a population of cells co-expressing Arc and NeuN in the ILPFC (9.84%) after fear extinction training.

**Suppl. Fig. 2.**
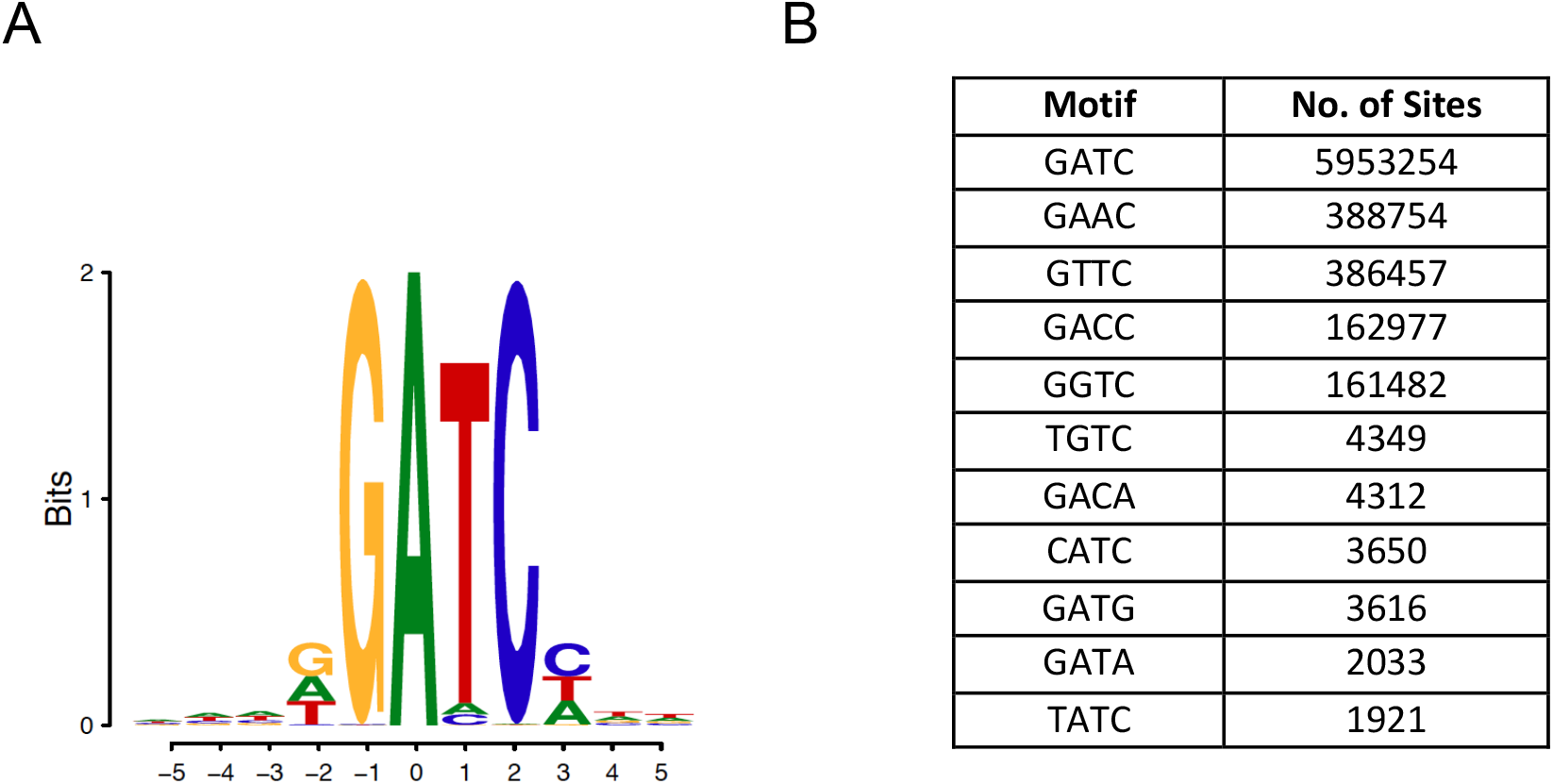
Motifs of m6dA sites in the mouse neuronal genome identified by DpnI-seq. (A) *De novo* motif result generated by HOMER Motif Analysis. The height represents the significance of enrichment of each nucleotide at that position. (B) List of all motifs that were identified by DpnI-seq followed by number of sites.

**Suppl. Fig. 3.**
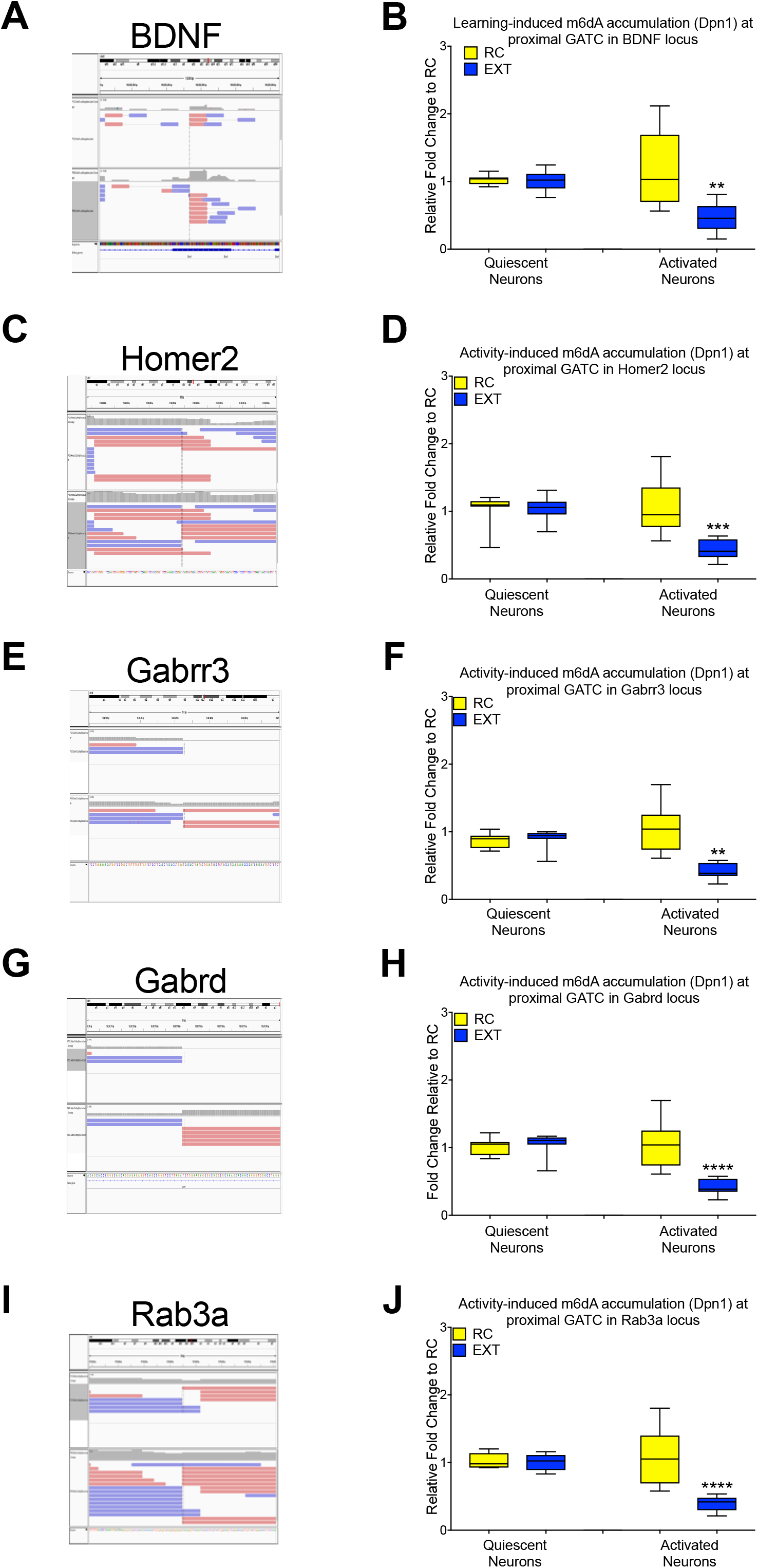
Learning-induced accumulation of m6dA only occurs in activated neurons. (A, C, E G and I) Raw reads image of DpnI-seq data from IGV browser. Dpnl-qPCR results reveal cell type specific accumulation of m6dA at (B) BDNF P4 proximal GATC site (two-way ANOVA F_1,28_ = 11.52, p<.01; Dunnett’s posthoc test: retention control (RC) vs. extinction (EXT) within activated neurons, **p<.01), (D) Homer2 (two-way ANOVA F_1,28_ = 11.38, p<.01; Dunnett’s posthoc test: retention control (RC) vs. extinction (EXT) within activated neurons, ***p<.001); (E) Gabrr3 (two-way ANOVA F_1,27_ = 6.703, p<.05; Dunnett’s posthoc test: RC vs. EXT within activated neurons, **p<.01); (H) Gabrd (two-way ANOVA F_1,27_ = 14.21, p<.001; Dunnett’s posthoc test: RC vs. EXT within activated neurons, ****p<.0001) and (J) Rab3a (two-way ANOVA F_1,28_ = 17.21, p<.001; Dunnett’s posthoc test: RC vs. EXT within activated neurons, ****p<.0001). (All n=7-8/group, Error bars represent S.E.M.).

**Suppl. Fig. 4.**
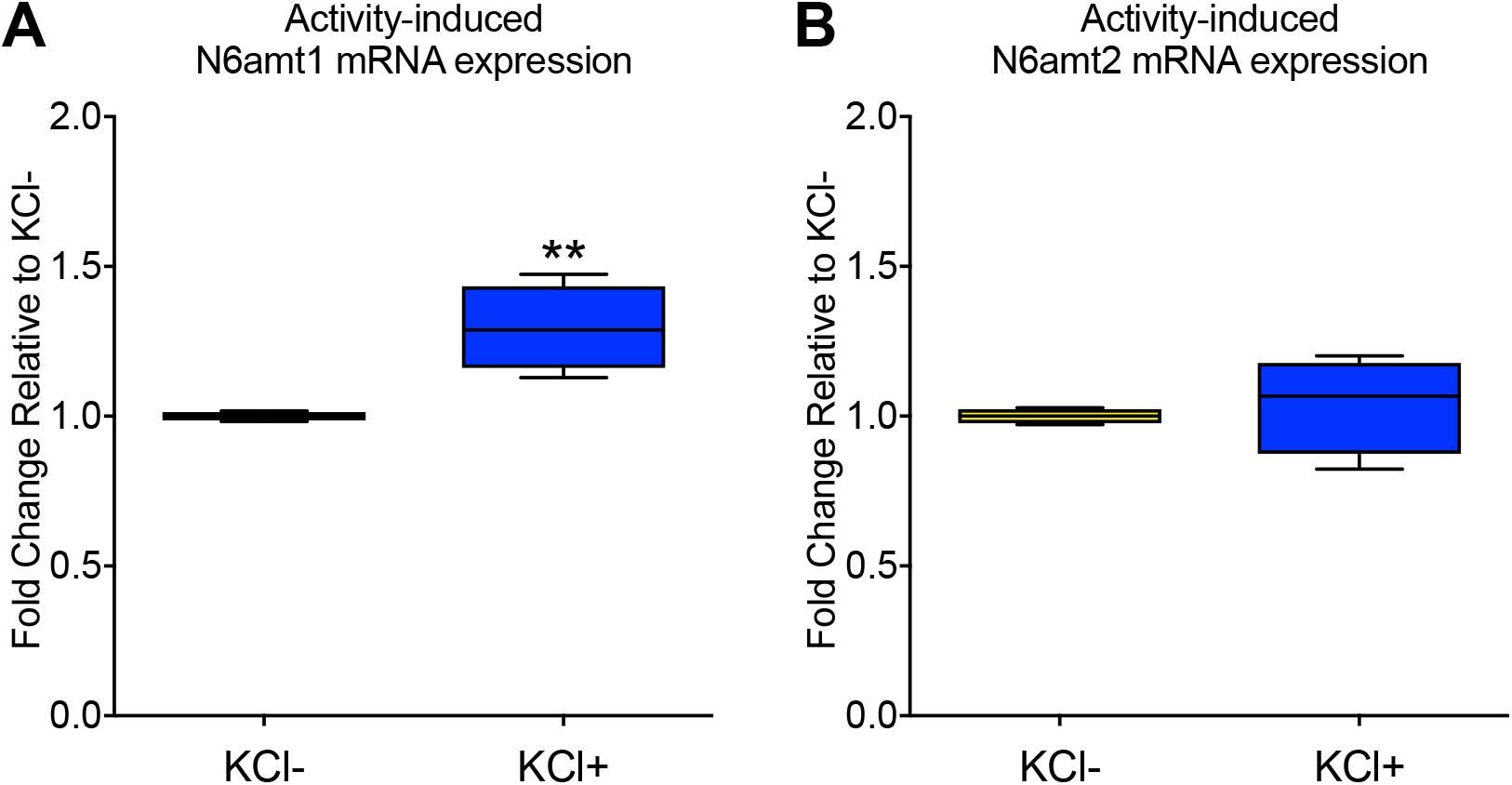
N6amt1 mRNA expression is extinction learning induced in primary cortical neurons. (A) Activity-induced N6amt1 mRNA expression in primary cortical neurons, *in vitro* (two-tailed, unpaired student’s t test, t=4.411, df=6,**p<.01). (B) No effect of neuronal stimulation on N6amt2 mRNA expression.

**Suppl. Fig. 5.**
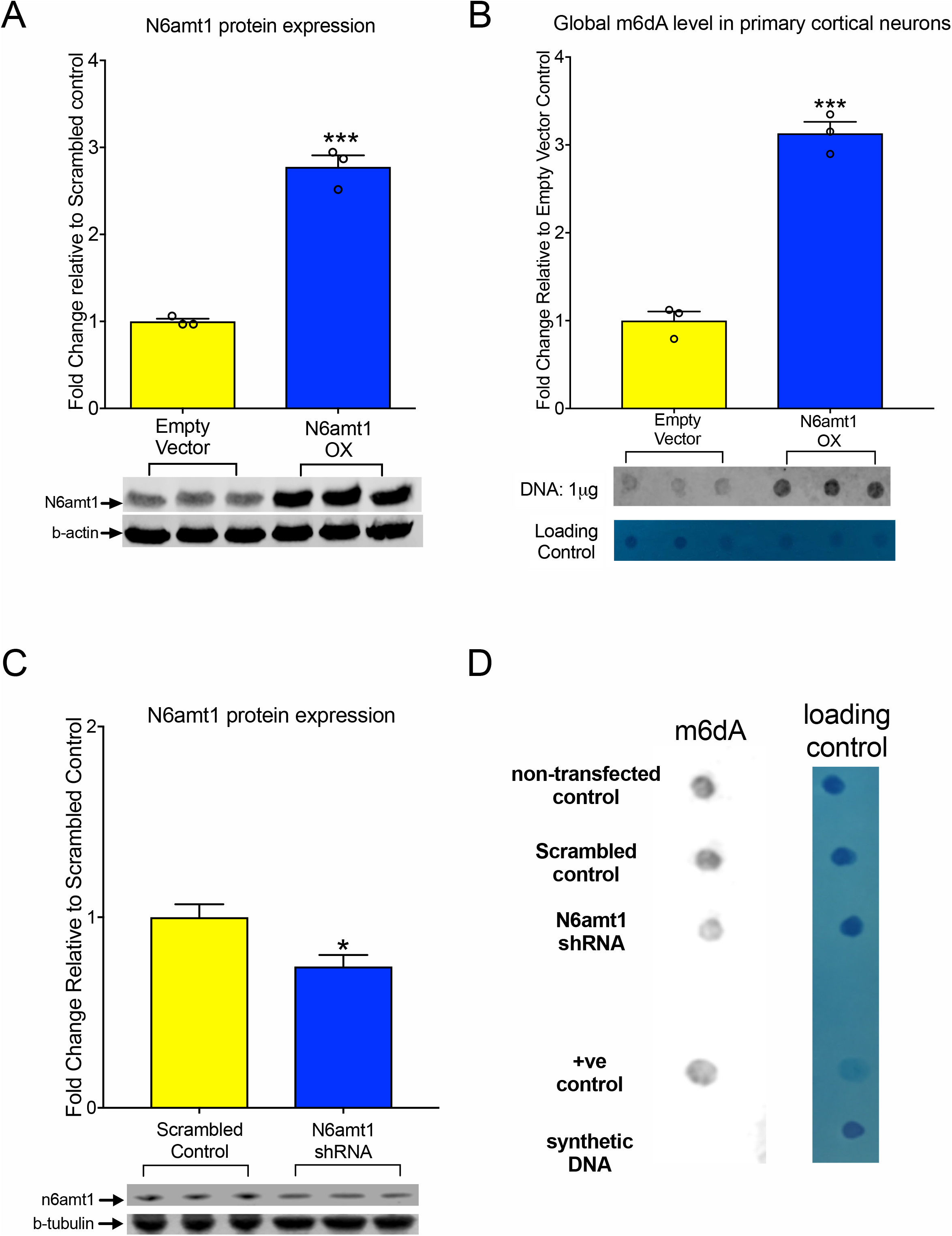
N6amt1 overexpression leads to a global increase in m6dA, and n6amt1 knockdown reduce global level of m6dA, *in vitro*. (A) N6amt1 protein levels in HEK293t cells shows N6amt1 overexpression construct can increased N6amt1 expression two-tailed, unpaired student’s t test, t=13.08, df=4,***p<.001). (B) N6amt1 over expression leads to an increase in the global level of m6dA in primary cortical neurons (two-tailed, unpaired student’s t test, t=12.75, df=4,***p<.001). (All n=3/group, Error bars represent S.E.M.). (C) N6amt1 shRNA knockdown leads to an reduction of protein level in primary cortical neurons (two-tailed, unpaired student’s t test, t=2.845, df=4,*p<.05), which leads to (D) reduction of global level of m6dA.

**Suppl. Fig. 6.**
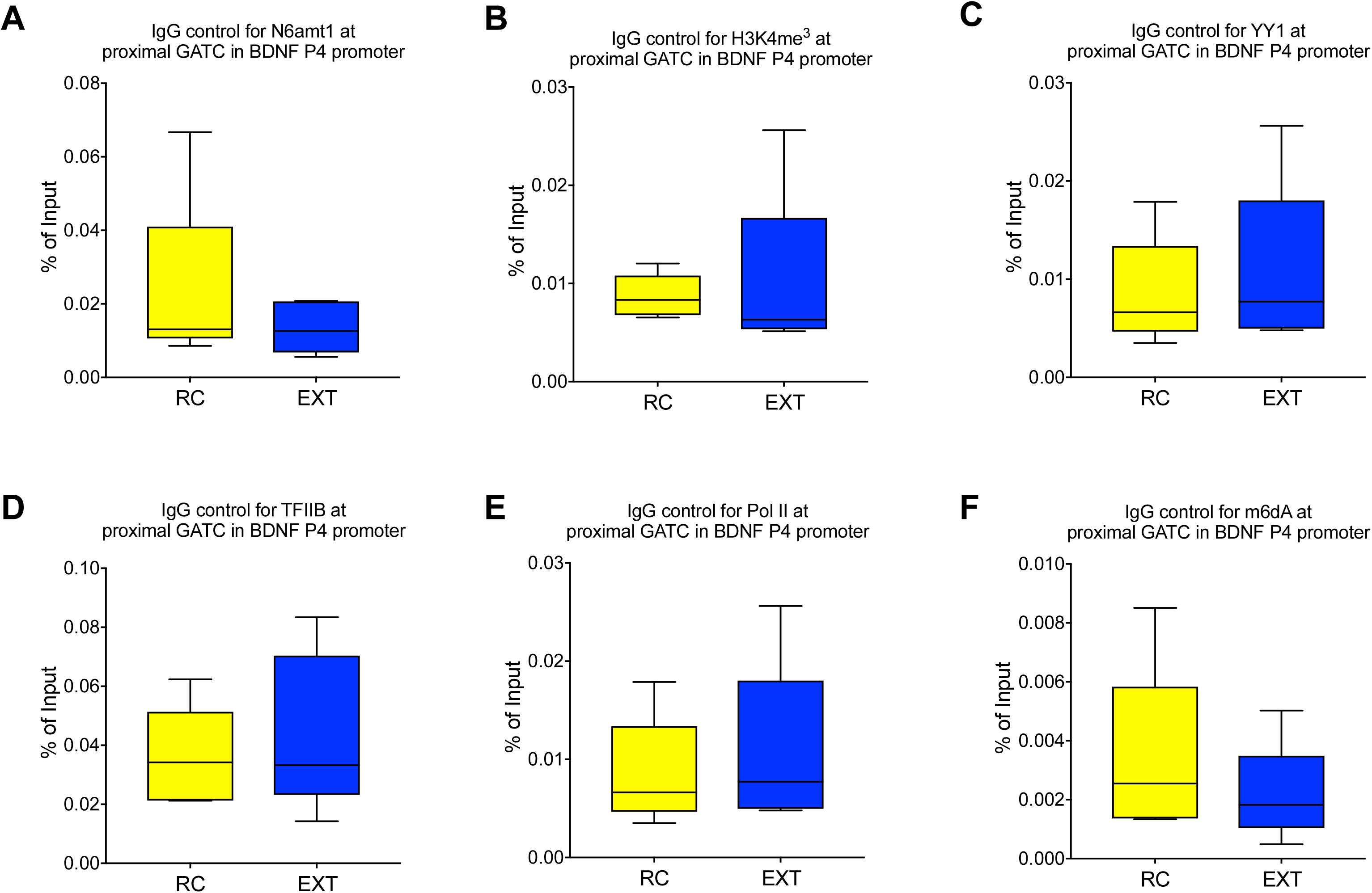
ChIP-qPCR with IgG controls of selected antibodies against N6amt1, H3K4me^3^, YY1, PolII and m6dA. There is no significant difference between retention control (RC) and extinction (EXT) at the proximal GATC site in the BDNF P4 promoter with (A) IgG control for N6amt1, (B) IgG control for H3K4me^3^, (C) IgG control for YY1, (D) IgG control for TFIIB, (E) IgG control for PolII and (F) IgG control for m6dA. (All n=5/group, Error bars represent S.E.M.).

**Suppl. Fig. 7.**
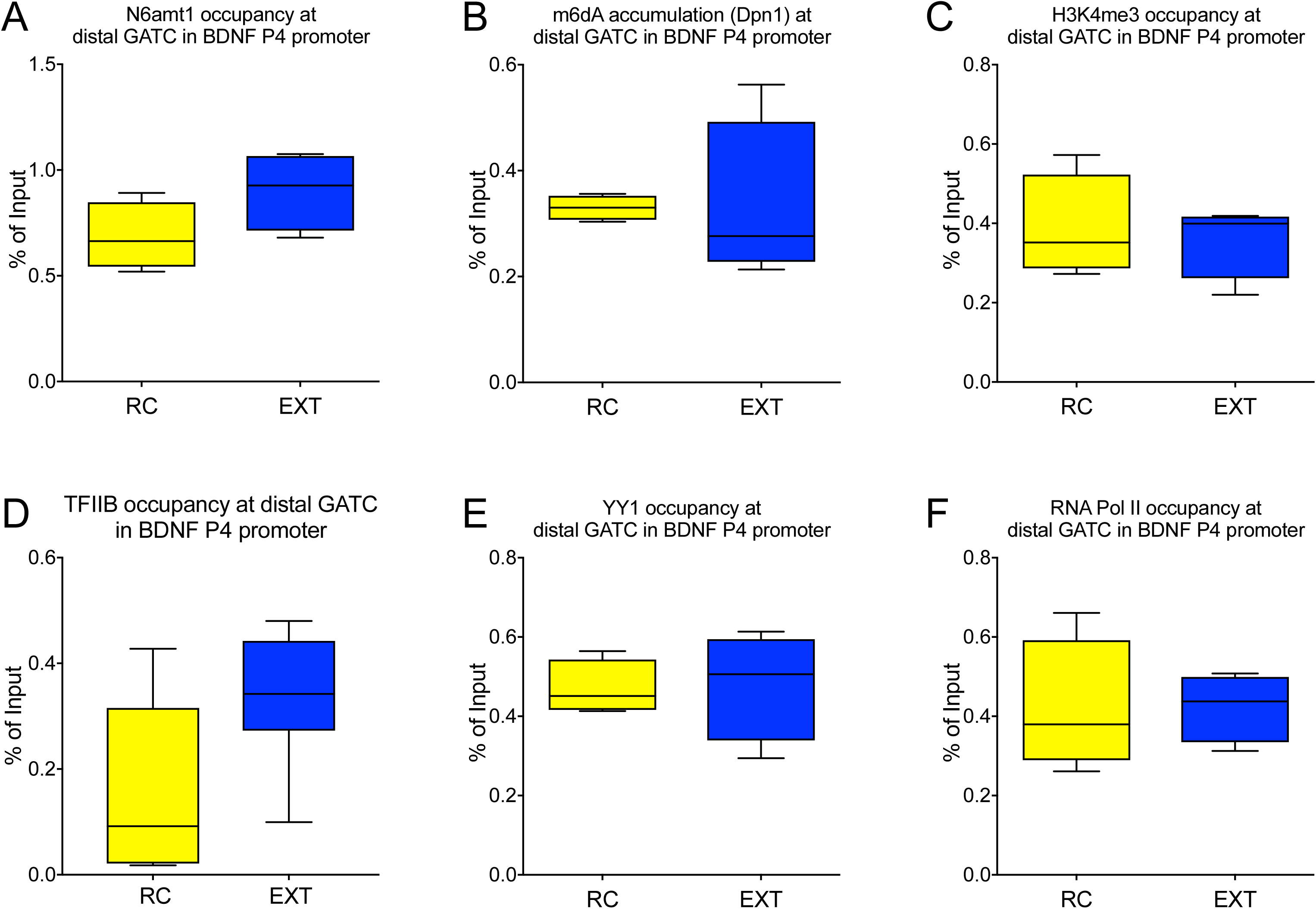
N6amt1-mediated and m6dA-related changes in chromatin and transcriptional machinery do not occur at a distal GATC sequence in the BDNF P4 promoter in ILPFC after learning. Extinction induced effect on (A) N6amt1 occupancy (B) the deposition of m6dA, or (C-F) the presence of open chromatin structure, H3K4me^3^, YY1 and RNA Pol II at the distal GATC site within the BDNF P4 promoter. (All n=4/group, Error bars represent S.E.M.).

**Suppl. Fig. 8.**
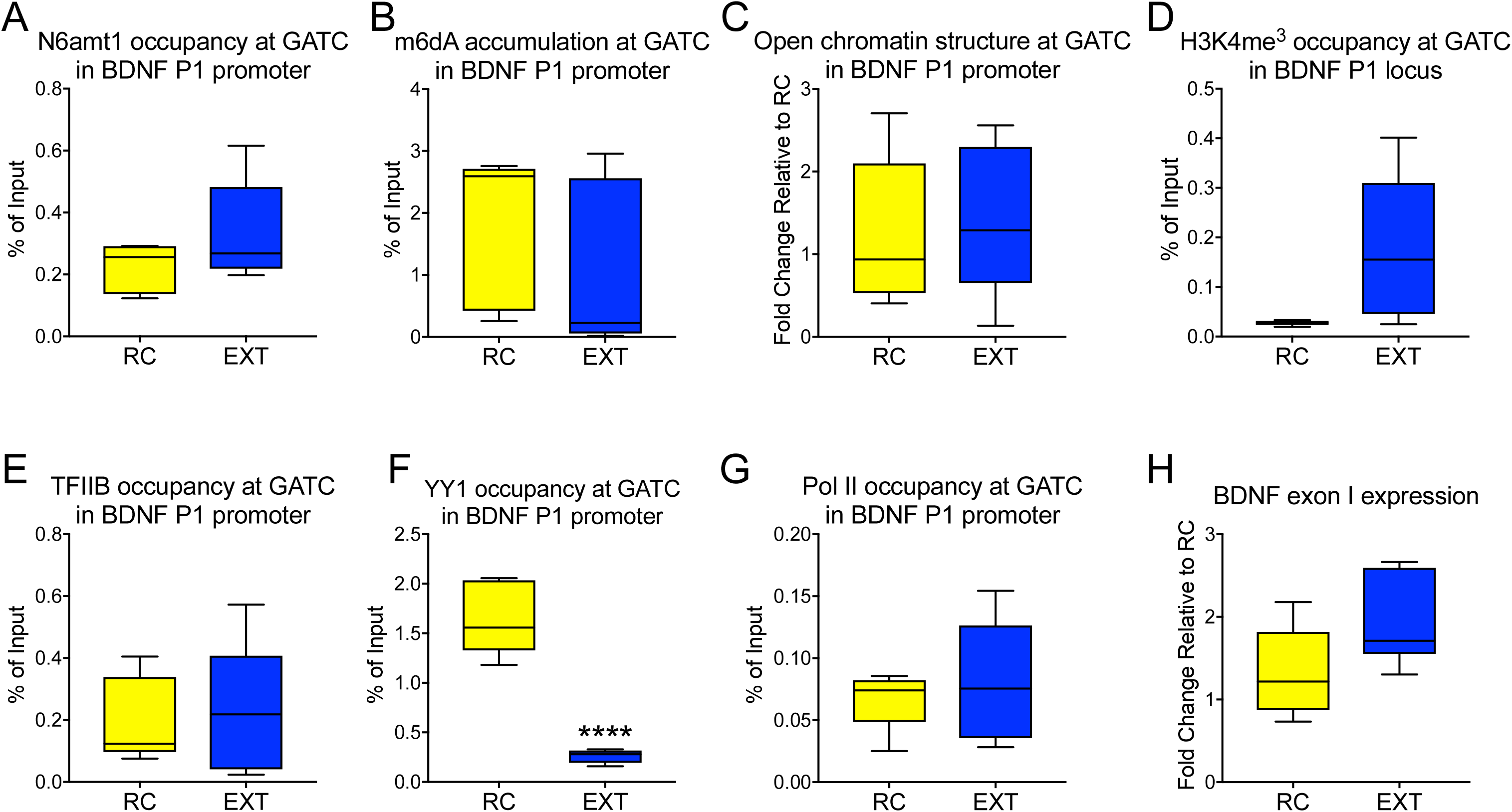
N6amt1-mediated and m6dA-related changes in chromatin and transcriptional machinery do not occur at the BDNF P1 promoter in ILPFC after learning. Extinction induced effect on (A) N6amt1 occupancy (B) the deposition of m6dA, or (C) the presence of open chromatin structure, (D) H3K4me^3^ and (E) TFIIB; (F) observed a reduction on YY1 occupancy (two-tailed, unpaired student’s t test, t=8.24, df=8, ****p<.0001). Also, no significant induction is detected on (G) RNA Pol II at the distal GATC site within the BDNF P4 promoter. (H) BDNF exon I expression (All n=4/group, Error bars represent S.E.M.).

**Suppl. Fig. 10.**
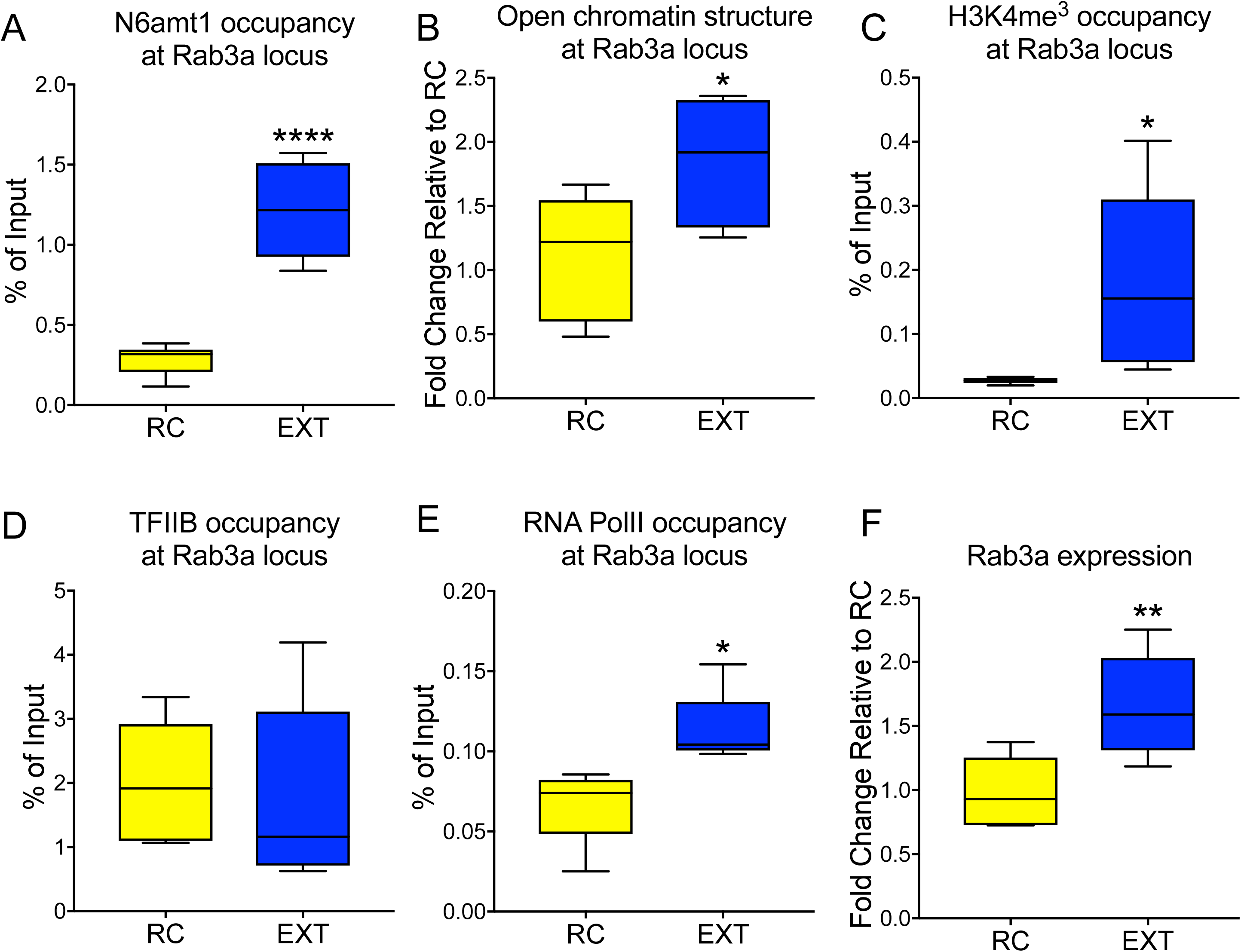
Extinction learning induces N6amt1 occupancy and active chromatin structures that are associated with increased Rab3a mRNA expression. Extinction training leads to (A) an increase in N6amt1 (two-tailed, unpaired student’s t test, t=7.252, df=10, ****p<.0001), (B) increased open chromatin structure (two-tailed, unpaired student’s t test, t=2.372, df=8, *p<.05), (C) increased occupancy of the chromatin mark H3K4^me3^ (two-tailed, unpaired student’s t test, t=2.334, df=8, *p<.05), however, no significant induction of (D) YY1 recruitment. On the other hand, (G) an increase in the presence of Pol II was observed (two-tailed, unpaired student’s t test, t=3.111, df=8, *p<.05), and (H) a correlated increase in the induction of Rab3a mRNA expression (two-tailed, unpaired student’s t test, t=3.471, df=10, **p<.01). (All n=5-6/group, Error bars represent S.E.M.).

**Suppl. Fig. 11.**
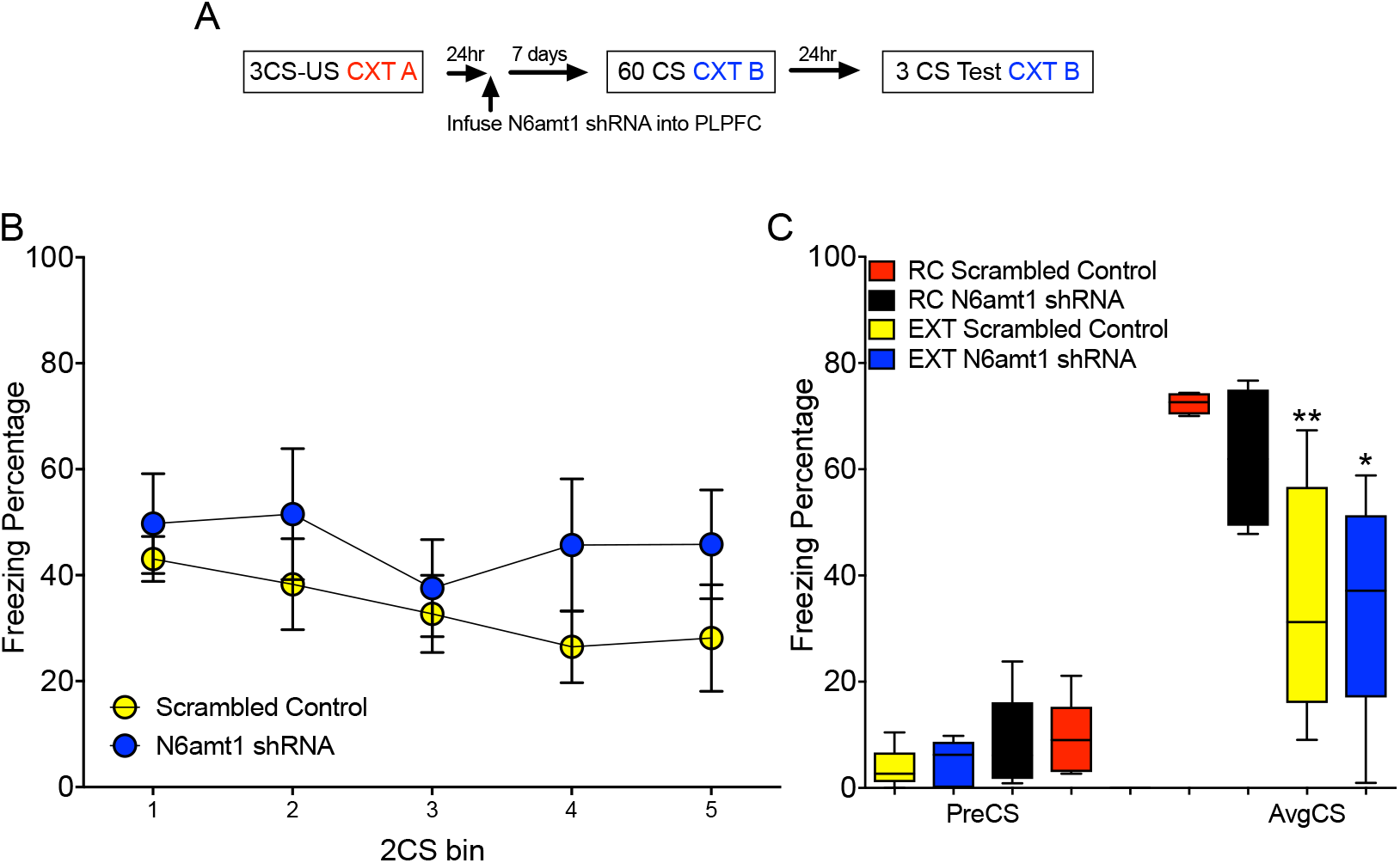
N6amt1 knockdown in the PLPFC has no effect on the formation of fear extinction memory. (A) The schema describes the behavioral test protocol. (B,C) knockdown of N6amt1 in PLPFC has no effect on the formation of extinction formation (one-way ANOVA F_8,34_ = 16.71, p<.0001; Tukey’s posthoc test: scrambled control RC vs. scrambled control EXT, **p<.01; N6amt1 shRNA RC vs. N6amt1 shRNA EXT, *p<.05). (All n=5-6/group, Error bars represent S.E.M.).

## Materials and Methods

### Mice

Male C57BL/6 mice (10-12 weeks old) were housed four per cage, maintained on a 12hr light/dark time schedule, and allowed free access to food and water. All testing took place during the light phase in red-light-illuminated testing rooms following protocols approved by the Institutional Animal Care and Use Committee of the University of California, Irvine and by the Animal Ethics Committee of The University of Queensland. Animal experiments were carried out in accordance with the Australian Code for the Care and Use of Animals for Scientific Purposes (8th edition, revised 2013).

### DNA/RNA extraction

Tissue derived from the ILPFC of retention control (RC) or extinction (EXT) trained mice was homogenized by Dounce tissue grinder in 500 μl cold 1X PBS (Gibco). 400 μl of homogenate was used for DNA extraction, and 100 μl was used for RNA extraction. DNA extraction was carried out using DNeasy Blood & Tissue Kit (Qiagen) with RNAse A (5 prime), RNAse H and RNAse T1 treatment (Invitrogen), and RNA was extracted using Trizol reagent (Invitrogen). Both extraction protocols were conducted according to the manufacturer’s instructions. The concentration of DNA and RNA was measured by Qubit assay (Invitrogen).

### LC-MS/MS

Genomic DNA was enzymatically hydrolyzed to deoxynucleosides by the addition of benzonase (25 U, Santa Cruz Biotech), nuclease P1 (0.1 U, Sigma-Aldrich), and alkaline phosphatase from *E.coli* (0.1 U, Sigma-Aldrich) in 10 mM ammonium acetate pH 6.0, 1 mM MgCl_2_, and 0.1 mM erythro-9-(2-hydroxy-3-nonyl) adenine. After 40 min incubation at 40 °C 3 volumes of acetonitrile was added to the samples and centrifuged (16,000 g, 30 min, 4 °C). The supernatants were dried and dissolved in 50 μl 5% methanol in water (v/v) for LCMS/MS analysis of modified and unmodified nucleosides. Chromatographic separation was performed on a Shimadzu Prominence HPLC system, for m6dA and unmodified deoxynucleosides by means of an Ascentis Express F5 150 x 2.1 mm i.d. (2.7 μm) column equipped with an Ascentis Express F5 12.5 × 2.1 mm i.d. (2.7 μm) guard column (Sigma-Aldrich). The mobile phase consisted of water and methanol (both containing 0.1 % formic acid), for m6dA, starting with a 4-min gradient of 5-50 % methanol, followed by 6 min re-equilibration with 5% methanol, and for unmodified deoxynucleosides maintaining isocratically with 30% methanol. The mobile phase consisted of 5 mM acetic acid and methanol, starting with a 3.5-min gradient of 5-70% methanol, followed by 4 min re-equilibration with 5% methanol. Mass spectrometry detection was performed using an API5500 triple quadrupole (AB Sciex) operating in positive electrospray ionization mode for m6dA and unmodified deoxynucleosides, or negative mode for m6dA.

### Dot blot

100 ng of total DNA was diluted with 0.1 N NaOH (Sigma) into 2 μl and spotted 2 μl on a nitrocellulose membrane (BioRad). DNA was spotted on the membrane and followed by 10 min incubation at room temperature. DNA was hybridized to membrane using 15 min incubation at 80 °C. The membrane was blocked in blocking buffer (Licor) for 60 min. The membrane was then incubated with 1:1,000 dilution of m6A (Active Motif) at 4 °C overnight. After three rounds of washes with 1X TBST, the membrane was incubated with 1:15,000 goat anti-rabbit secondary antibody conjugated with AlexaFluor 680 (LiCOR). The membranes were then washed with 1X TBST and imaged, and the intensity score of each dot was analyzed by the Odyssey Fc system and normalized to background.

### qRT-PCR

500 ng RNA was used for cDNA synthesis using the PrimeScript Reverse Transcription Kit (Takara). Quantitative PCR was performed on a RotorGeneQ (Qiagen) cycler with SYBR-Green Master mix (Qiagen) using primers for target genes and for beta-actin as an internal control (Suppl. Table 1). All transcript levels were normalized to beta-actin mRNA using the _δδ_CT method, and each PCR reaction was run in duplicate for each sample and repeated at least twice.

### DNA shearing

DNA and chromatin was sheared using m220 Ultra-sonicator (Covaris) with an average size about 300 bp. The program set as Peak Power: 50, Duty Factor: 20, Cycle/Burst: 200, Duration: 75 sec and Temperature: 18°C to 22°C.

### Chromatin immunoprecipitation

Chromatin immunoprecipitation (ChIP) was performed following modification of the Invitrogen ChIP kit protocol. Tissue was fixed in 1% formaldehyde and cross-linked cell lysates were sheared by Covaris in 1% SDS lysis buffer to generate chromatin fragments with an average length of 300bp by using Peak Power: 75, Duty Factor: 2, Cycle/Burst: 200, Duration: 900 Secs and temperature: between 5 °C to 9 °C. The chromatin was then immunoprecipitated using previously validated antibodies for H3K4me^3 1^, YY1 ^2^, TFIIB^3^ and RNA Pol II ^4^. Also, an equivalent amount of normal rabbit IgG or mouse IgG (Santa Cruz) was used for non-specificity control. Antibody and sample mixtures were then incubated overnight at 4°C. Protein-DNA-antibody complexes were precipitated with protein G-magnetic beads (Invitrogen) for 1hr at 4°C, followed by three washes in low salt buffer, and three washes in high salt buffer. The precipitated protein-DNA complexes were eluted from the antibody with 1% SDS and 0.1 M NaHCO_3_, then incubated 4hr at 60°C in 200 mM NaCl to reverse the formaldehyde cross-link. Following proteinase K digestion, phenol-chloroform extraction, and ethanol precipitation, samples were subjected to qPCR using primers specific for 200 bp segments corresponding to the target regions.

### Dpnl-Seq

Frozen ILPFC tissues were homogenized and fixed with 1% methanol free PFA (Thermo) at room temperature for 5 mins. Final concentration of 0.125 mM of glycine was then added to stop fixation. The cells were then washed with 1X cold PBS for three times and the cell suspension treated with DNaseI (Thermo) for 15 mins at 4 °C followed by 1ml of 1X cold PBS wash. The cell suspension was then blocked by using FACS blocking buffer (1X BSA, 1X normal Goat Serum and 1% TritonX) for 15 min at 4 °C with end-to-end rotation. After 15 min., the cell suspension was incubated with 1:150 dilution of preconjugated Arc antibody (Bioss) and 1:300 dilution of preconjugated NeuN antibody (Bioss) at 4°C for 1hr with an end to end rotation. Following incubation, two rounds of 1ml 1X cold PBS washes were applied. Then, the cell pellets were resuspended with 500μl of 1X cold PBS, and 1:2000 DAPI was added. FACS was performed on a BD FACSAriaII (BD Science). DNA was extracted from FACS sorted cells by phenol/chloroform. -seq library preparation was modified from a previously published protocol ^5^ and samples were sequenced on a HiSeq4000.

### FACS sorted RNA-seq

FACS sorting was performed as previously described. RNA was isolated from sorted cells by Arcturus PicoPure RNA isolation kit (Applied Biosystems) following the manufacturers protocol. 5ng of total RNA per sample and SMRTer Stranded Total RNA-seq Kit v2 Pico Input Mammalian Components kit (Clontech) was used for RNA-seq library preparation following the manufacturers protocol and samples sequenced on a HiSeq2500 within the QBI genome sequencing facility.

### N6amt1 ChIP-seq

N6amt1 ChIP was performed as previously described. After chromatin immunoprecipitation (using 4μg antibody), DNA was extracted by using the DNA clean & Concentrator kit (Zymo Research). 10ng of enriched DNA per sample and KAPA DNA HyperPrep kit (Roche) was used for ChIP-seq library preparation following the manufacturers protocol. The sequencing run was performed on the Illumina Nextseq platform.

### Dpnl-seq Data analysis

Illumina pair-end sequencing data was aligned to the mouse reference genome (mm10) using BWA (v0.6.2) ^6^. Samtools (v0.1.17) ^7^ was then used to convert “SAM” files to “BAM” files, sort and index the “BAM” files, and remove duplicate reads. Reads with low mapping quality (<20) or reads that were not properly paired-end aligned to the reference genome were excluded from the downstream analysis. These steps ensure that only high-quality alignments were used for the analysis of DpnI cleavage sites (Suppl. Table 2). After alignment, we applied a similar approach that infers potential DpnI cleavage sites based on the position of 5’ends as described in a previous study ^5^. Briefly, a binominal distribution model was assumed that each read could be randomly sheared and aligned to the genome with a probability *p* = = 1/gs (gs = genome size) or cleaved by DpnI. For each individual sample, let *n* be the total number of reads. The *P* value of each genomic locus supported by *x* number of reads was calculated as 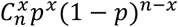. Bonferroni correction was then applied for multiple testing correction. A genomic locus was determined as a real DpnI cleavage site if it satisfies the following criteria: i) the corrected *P* value < 0.01 in at least 2 of the 3 biological replicates in one condition or both conditions, and ii) the locus is not in the mm10 empirical blacklists identified by the ENCODE consortium ^8^.

The detected DpnI cleavage sites in each condition as well as merged data were used for motif analysis, separately. A Dpnl cleavaged “GATC” site was determined as a differentially methylated site between RC and EXT conditions if it satisfies i) the DpnI cleavage site are supported by at least two biological replicates in one condition (e.g. condition A) but at most one replicate in the other condition (e.g. condition B), ii) all three biological replicates in condition A should have 5’ end(s) supporting the DpnI site, and iii) the number of 5’end supported reads in condition A is at least two-folds more than that in condition B. Genes with differentially methylated GATC sites near the TSS region (+/− 500 bp) were parsed for GO enrichment analysis using DAVID (version 6.8) ^9,10^.

### N6amt1 ChIP-Seq data analysis

We performed the paired-end reads alignment and filter using the same analysis workflow as described in Dpnl-Seq data analysis. After removed duplicate reads, low mapping quality reads, and not properly paired-end aligned reads, MACS2 (version 2.1.1.20160309) was used to call peaks for each sample with the parameter setting “callpeak -t SAMPLE -c INPUT -f BAMPE --keep-dup=all -g mm -p 0.05 -B”. Peak summits identified by MACS2 from all samples were collected to generate a list of potential binding sites. Custom PERL script was then applied to parse the number of fragments (hereafter referred as counts) that cover the peak summit in each sample. Each pair of properly paired-end aligned reads covering the peak summit represents one count. The total counts in each sample were normalised to 20 million before comparison among samples. The potential binding sites were kept if they met all of the following conditions: i) the sites were not located in the mm10 empirical blacklists, and ii) the normalized counts in all three biological replicates in one group were larger than that in its input sample, and the normalized counts in at least 2 replicates were more than 2-folds larger than its normalized input count.

### RNA-Seq data analysis

Illumina paired-end reads were aligned to the mouse reference genome (mm10) using HISAT2 (version 2.0.5), with the parameter setting of “--no-unal --fr --known-splicesite-infile mm10_splicesites.txt”. The “htseq-count” script in HTSeq package (v0.7.1) (http://www-huber.embl.de/HTSeq) was used to quantitate the gene expression level by generating a raw count table for each sample. Based on these raw count tables, edgeR (version 3.16.5) was adopted to perform the differential expression analysis between groups. EdgeR used a trimmed mean of M-values to compute scale factors for library size normalization. Genes with counts per million (CPM) > 1 in at least 3 samples were kept for downstream analysis. We applied the quantile-adjusted conditional maximum likelihood (qCML) method to estimate dispersions and the quasi-likelihood F-test to determine differential expression. Differentially expressed genes between two groups were identified when FDR < 0.05. Gene ontology enrichment analysis for differentially expressed genes was performed using the functional annotation tool in DAVID Bioinformatics Resources (version 6.8).

### Determine distance from m6dA sites to N6amt1 sites

For each N6amt1 binding site (peak summit) locating in TSS +/− 500bp regions, we searched its nearby m6A sites and extracted the distance between the peak summit and its closest m6A site. The distribution plot shows that most N6amt1 binding sites have a nearby m6A site within 500bp, with 0-200 being the most abundant.

### m6dA MeDIP-DIP

1 μg of genomic DNA was diluted to 130 μl ultrapure water (Invitrogen) and sheared with an average size about 300bp prior to the capture. m6dA captured was performed using a m6dA antibody (Active Motif) to capture m6dA enriched genomic regions. The procedure was adapted from manufactory’s protocol for Methyl DNA immunoprecipitation (Active motif). 500 ng of sheared DNA and 4 μg of m6dA antibody was used for each immunoprecipitation reaction and all selected targets (GATC site proximal BDNF P4: Chr2: 109692436-109692774; distal GATC site: Chr2: 109691953-109692103) were normalized to input DNA and then to their own controls by using the _δδ_CT method, and each qPCR reaction was run in duplicate for each sample and repeated at least twice.

### Dpnl-qPCR

300 ng of sheared DNA was treated with 200 units of Dpnl (NEB) for 16 hours at 37°C and followed by heat inactivation using 80 °C for 20 min. Treated DNA was then used in qPCR reactions. All selected targets were normalized to its own untreated control by using the _δδ_CT method, and each PCR reaction was run in duplicate for each sample and repeated at least twice. A schema is included in figure 1B.

### FAIRE (Formaldehyde-Assisted Isolation of Regulatory Elements)-qPCR

The procedure was adapted from a published protocol^11 12^ ILPFC tissues were homogenized within 500ul of 1X cold PBS by duncer (10 strocks). Molecular grade 16% of formaldehyde (Thermofisher) was added directly to the cell suspension at room temperature (22-25°C) to a final concentration 1% and incubate for 5min, Glycine was added to a final concentration of 125mM for 5min at room temperature to stop fixation. Two rounds of 1x cold PBS wash was performed to wash off remind formaldehyde. The cells were collected by centrifugation at 2000rpm for 4mins and store at −80 °C. Fixed cell pellets was then treated with chIP lysis buffer as described above and samples were then sonicated by using Covaris to generate chromatin fragments with an average length of 300bp by using Peak Power: 75, Duty Factor: 2, Cycle/Burst: 200, Duration: 900 Secs and temperature: between 5 °C to 9 °C. Cellular debris was cleared by spinning at 15,000 rpm for 10mins at 4°C. DNA was isolated by adding an equal volume of an equal volume of phenol-chloroform (Sigma) and followed by vertexing and spinning at 15,000 rpm 15mins at 4°C. The aqueous phase was isolated and store into another 1.5ml microcentrifuge tube. An additional 500ul of TE buffer was added to the organic phase, vortexed and centrifuged again at 15,000rpm for 15mins at 4°C. The aqueous phase was isolated and combined with the first aqueous fraction. Another phenol-chloroform extraction was performed on the pooled aqueous fractions to ensure that all protein was removed. The DNA was isolated by followed previous described DNA extraction procedures within chIP protocol. Input DNA isolation were performed same as previous described. qPCR was performed using SYBR green master mix (Qiagen) on a Rotorgene platform (Qiagen). Relative enrichment of each target in the FAIRE-treated DNA was calculated based on untreated input DNA.

### N6amt1 knockdown constructs

Lentiviral plasmids were generated by inserting either N6amt1, N6amt2 shRNA or scrambled control fragments (supplementary table 1) immediately downstream of the human H1 promoter in a modified FG12 vector (FG12H1, derived from the FG12 vector originally provided by David Baltimore, CalTech) as previously described ^13^. Lentivirus was prepared and maintained according to protocols approved by the Institutional Biosafety Committee at the University of California Irvine.

### N6amt1 over-expression lentiviral constructs

Lentiviral plasmids were generated by inserting a full mouse N6amt1 transcript cDNA with GFP from FUGW (addgene) backbone. First, a full length of n6amt1 cDNA was PCR amplified from N6amt1 (Myc-DDK-tagged) construct (Origene cDNA clone#MR227618). The forward primer is 5’-ATTCGTCGACTGGATCCGGT. The reverse primer is 5’-CCGAATTCGGCCGGCCGTTTAAACCTT. The PCR amplified fragments were inserted immediately downstream of the Ubiquitin C promoter of a lentiviral vector, FUIGW-K1. The original FUGW vector was used as an Empty vector control. Then, either N6amt1 over expression or control plasmid was co-transfected with lentiviral helper plasmids (pMDL, pVSVG and pREV) into HEK 293T cells with ~80% confluence. 4 hours later, Sodium butyrate was added to stimulate viral production. After two days incubations at 37°C and 5% CO2, virus were collected by ultracentrifugation. The titer was measured with Lenti-X Gostix (Clontech).

### Titration of Virus

4×10^5 293T cells/well of a 6 well plate were plated the day before titration of virus. Next day, number of cells in each well were estimated by counting. Then, each virus was added 0.5, 1, 2 and 5ul per well and Incubated for 2-3 days. Percentage of cells expressing EGFP was used to calculate the virus titre by using following formula: % GFP positivie cells X 1/ml of virus added to well X number of cells in well prior to infection = Infectious units (IU)/ml X 10^3 = IU/ml. Only virus that reached over 1X10^8 IU/ml was used within this study.

### Cannulation surgery and lentiviral infusion

A double cannula (PlasticsOne) was implanted in the anterior posterior plane, +/− 30 degrees along the midline, into the ILPFC, a minimum of 3 days prior to viral infusion. The coordinates of the injection locations were centered at +1.80 mm in the anterior-posterior plane (AP), −2.7 mm in the dorsal-ventral plane (DV). For PLPFC cannulation surgery, the coordinates of the injection locations were centered at +1.80 mm in the anterior-posterior plane (AP), −2.0 mm in the dorsal-ventral plane (DV), and 0mm in the medial-lateral plane (ML). 1.0 ul of lentivirus was introduced bilaterally via 2 injections delivered within 48 hours. For knockdown experiments, mice were first fear conditioned, followed by 2 lentiviral infusions 24 hours post fear condition training, and after a one-week incubation; then, extinction trained. After training, viral spread was verified by immunohistochemistry according to a previously published protocol^14,15^

### Lentiviral knockdown and overexpression (Ox) of N6amt1, *in vitro*

1 μl of N6amt1 shRNA/OX or scrambled control/empty vector control lentivirus was applied to primary cortical neurons in a 6-well plate. After 7 days incubation, cells were harvested for RNA extraction.

### Behavioral Tests

Two contexts (A and B) were used for all behavioral fear testing. Both conditioning chambers (Coulbourn Instruments) had two transparent walls and two stainless steel walls with a steel grid floors (3.2 mm in diameter, 8 mm centers); however, the grid floors in context B were covered by flat white plastic non-transparent surface with two white LED lights to minimize context generalization. Individual digital cameras were mounted in the ceiling of each chamber and connected via a quad processor for automated scoring of freezing measurement program (FreezeFrame). Fear conditioning was performed in context A with spray of vinegar (10% distilled vinegar). Then, actual fear condition protocol was starting with 120 sec pre-fear conditioning incubation; then, followed by three pairing of a 120 sec, 80dB, 16kHZ pure tone conditioned stimulus (CS) co-terminating with a 1 sec (2 min intervals), 0.7 mA foot shock (US). Mice were counterbalanced into equivalent treatment groups based on freezing during the third training CS. For extinction, mice were exposed in context B with a stimulus light on and spray of Almond (10% Almond extracts and 10% ethanol). Mice allowed to be acclimated for 2 min, and then, extinction training comprised 60 non-reinforced 120 sec CS presentations (5-sec intervals). For the behavior control experiments, context exposure was performed for both fear condition and fear extinction training. Animal, inside 3CS-US or 60CS treatment, only exposed into either context A or B with equal times of mice spend there by fear condition or extinguished mice but were not exposed to any 3CS-US or 60CS. For the retention test, all mice were returned to context B and following a 2 min acclimation (used to minimize context generalization), freezing was assessed during three 120 sec CS presentations (120 sec intertribal interval). Memory was calculated as the percentage of time spent freezing during the tests.

### Behavioral Training (for tissue collection)

Naïve animals remained in their home cage until sacrifice. For the other groups, fear conditioning consisted of three pairing (120 sec inner-trial interval ITI) of a 120 sec, 80dB, 16kHZ pure tone conditioned stimulus (CS) Co-terminating with a 1 sec, 0.7 mA foot shock in context A. Mice were matched into equivalent treatment groups based on freezing during the third training CS. Context A exposure group spent an equivalent amount of time in context A without any CS and US. One day later, the fear-conditioned mice were brought to context B, where the extinction group (EXT) was presented with 60 CS presentations (5s ITI). The fear-conditioned without extinction (FC No-EXT) group spent an equivalent amount of time in context B without any CS presentations. Tissue was collected from both of these groups immediately after the end of either context B exposure (FC-No EXT) or extinction training (EXT).

### Primary cortical neuron, N2A and HEK cell culture

Cortical tissue was isolated from E15 mouse embryos in a sterile atmosphere. Tissue was dissociated by finely chopping, followed by gentle pipette to create single cell suspension. To prevent clumping of cells due to DNA from dead cells, tissue was treated with 2 unit/μl of DNase I. Cells finally went through the 40μm cell strainer (BD Falcon) and were plated onto 6 well plate coated with Poly-L- Ornithine (Sigma P2533) at a density of 1×10^6^ cells per well. The medium used was Neurobasal media (Gibco) containing B27 supplement (Gibco). 1X Glutamax (Gibco), and 1% Pen/Strep (Sigma). N2a cell was maintained in medium contains half DMEM, high glucose (GIBCO), half OptiMEM 1 (Gibco) with 5% serum and 1% Pen/Strep. HEK293t cell was maintained in medium contains DMEM, high glucose (Gibco) with 5% serum and 1% Pen/Strep (Gibco).

### Western blot

Protein samples were extracted by using NP40 solution followed manufactory’s protocol (Thermofisher) and protein concentration was determined by using Qubit protein detection Kit (Invitrogen) followed by manufactory’s protocol. Individual samples were run on a single 10 well gel or 12 well pre-made 4-12% gel (Thermofisher). Briefly, samples were prepared on ice (to final volume of 20 μl) and then vortexed and denatured for 10 min at 90 °C. Gels were run with 1X Tris buffered saline-Tween (TBS-T) and proteins transferred onto nitrocellulose membrane (Bio-rad). The membrane was blocked by blocking buffer (Licor) for 1hr at room temperature, washed with TBS-T for 5 min (3X), and incubated with 5ml of N6amt1 (1:250; Santa Cruz) and Beta-actin (1:500; Santa Cruz) or beta-tubulin (1:500; Santa Cruz) antibodies in blocking buffer (Licor) for overnight at 4 °C. The membranes were washed with TBS-T (3X), incubated for 1hr with anti-mouse secondary antibody (1: 15000; Li-Cor) and anti-rabbit secondary antibody (1:15000; Li-Cor) in blocking buffer (Li-Cor), and washed three times with TBST for 10 min (5X) and 20 min (1X). Optical density readings of the membrane were taken using a Li-Cor FX system followed by manufactory’s protocol.

### Statistical Analyses

In all cases where an unpaired t-test was employed, we opted for a one-tailed test with the a priori hypothesis that the accumulation of m6dA is permissive for gene expression and memory formation. Therefore, all experiments related to epigenetic and transcriptional machinery were hypothesized to show a positive correlation with m6dA, hence, a one-tailed test was employed. For behavioral analysis, freezing (the absence of all non-respiratory movement) was rated during all phases by automated digital analysis system, using a 5-sec instantaneous time sampling technique. The percentage of observations with freezing was calculated for each mouse, and data represent mean ± SEM freezing percentages for groups of mice during specified time bins. Total session means were analyzed with one-way ANOVA for the behavioral data in Figure 6 and Suppl. Fig.11. In experiments using viral manipulation and BDNF rescue, all data analysis was performed by two-way ANOVA for the data in Fig. 6 and 7, Suppl. Fig. 11. Dunnett’s posthoc tests were used where appropriate. For Dpnl-qPCR on FACS sorted samples in Suppl. Fig.3, two-way ANOVA followed by Dunnett’s posthoc tests.

## Acknowledgements

The authors gratefully acknowledge grant support from the NIH 5R01MH105398-TWB and PB; 5R01MH109588-03-RCS and TWB, the NHMRC GNT1033127-TWB, the Conselho Nacional de Desenvolvimento Científico e Tecnológico (CNPq-CsF-400850/2014-1-RGO), and the Research Council of Norway (FRIMEDBIO grant 32222-MB). XL has been supported by postgraduate scholarships from the University of Queensland and the ANZ Trustees Queensland and a postdoctoral fellowship from The University of Queensland. The authors would also like to thank Ms. Rowan Tweedale for helpful editing of the manuscript, and especially Sunil Gandhi for comments and lively discussion.

## References

1. Marshall, P. & Bredy, T. W. Cognitive neuroepigenetics: the next evolution in our understanding of the molecular mechanisms underlying learning and memory? NPJ Sci Learn 1, 16014 (2016).

2. Li, X. et al. Neocortical Tet3-mediated accumulation of 5-hydroxymethylcytosine promotes rapid behavioral adaptation. Proc. Natl. Acad. Sci. U.S.A. 111, 7120–7125 (2014).

3. Wei, W. et al. p300/CBP-associated factor selectively regulates the extinction of conditioned fear. J. Neurosci. 32, 11930–11941 (2012).

4. Miller, C. A., Campbell, S. L. & Sweatt, J. D. DNA methylation and histone acetylation work in concert to regulate memory formation and synaptic plasticity. Neurobiol Learn Mem 89, 599–603 (2008).

5. Baker-Andresen, D., Ratnu, V. S. & Bredy, T. W. Dynamic DNA methylation: a prime candidate for genomic metaplasticity and behavioral adaptation. Trends in Neurosciences 36, 3–13 (2013).

6. Gapp, K., Woldemichael, B. T., Bohacek, J. & Mansuy, I. M. Epigenetic regulation in neurodevelopment and neurodegenerative diseases. Neuroscience 264, 99–111 (2014).

7. Korlach, J. & Turner, S. W. Going beyond five bases in DNA sequencing. Curr. Opin. Struct. Biol. 22, 251–261 (2012).

8. Lister, R. et al. Global epigenomic reconfiguration during mammalian brain development. Science 341, 1237905–1237905 (2013).

9. Guo, J. U., Su, Y., Zhong, C., Ming, G.-L. & Song, H. Hydroxylation of 5-Methylcytosine by TET1 Promotes Active DNA Demethylation in the Adult Brain. Cell 145, 423–434 (2011).

10. Shen, L. et al. Tet3 and DNA replication mediate demethylation of both the maternal and paternal genomes in mouse zygotes. Cell Stem Cell 15, 459–470 (2014).

11. Khare, T. et al. 5-hmC in the brain is abundant in synaptic genes and shows differences at the exonintron boundary. Nat. Struct. Mol. Biol. 19, 1037–1043 (2012).

12. Miller, C. A. et al. Cortical DNA methylation maintains remote memory. Nature Neuroscience 13, 664–666 (2010).

13. Vanyushin, B. F., Mazin, A. L., Vasilyev, V. K. & Belozersky, A. N. The content of 5-methylcytosine in animal DNA: the species and tissue specificity. Biochim. Biophys. Acta 299, 397–403 (1973).

14. Iyer, L. M., Zhang, D. & Aravind, L. Adenine methylation in eukaryotes: Apprehending the complex evolutionary history and functional potential of an epigenetic modification. BioEssays 38, 27–40 (2016).

15. Hattman, S., Kenny, C., Berger, L. & Pratt, K. Comparative study of DNA methylation in three unicellular eucaryotes. J. Bacteriol. 135, 1156–1157 (1978).

16. Hattman, S. DNA-[Adenine] Methylation in Lower Eukaryotes. Biochemistry (Moscow) 70, 550–558 (2005).

17. Fu, Y. et al. N6-Methyldeoxyadenosine Marks Active Transcription Start Sites in Chlamydomonas. Cell 161, 879–892 (2015).

18. Zhang, G. et al. N6-Methyladenine DNA Modification in Drosophila. Cell 161, 893–906 (2015).

19. Greer, E. L. et al. DNA Methylation on N6-Adenine in C. elegans. Cell 161, 868–878 (2015).

20. Wu, T. P. et al. DNA methylation on N6-adenine in mammalian embryonic stem cells. Nature 532, 329–333 (2016).

21. Yao, B. et al. DNA N6-methyladenine is dynamically regulated in the mouse brain following environmental stress. Nat Comms 8, 1122 (2017).

22. Xiao, C.-L. et al. N6-Methyladenine DNA Modification in the Human Genome. Mol. Cell 71, 306–318.e7 (2018).

23. Usheva, A. & Shenk, T. TATA-binding protein-independent initiation: YY1, TFIIB, and RNA polymerase II direct basal transcription on supercoiled template DNA. Cell 76, 1115–1121 (1994).

24. Bredy, T. W. et al. Histone modifications around individual BDNF gene promoters in prefrontal cortex are associated with extinction of conditioned fear. Learn. Mem. 14, 268–276 (2007).

25. Liu, J. et al. Abundant DNA 6mA methylation during early embryogenesis of zebrafish and pig. Nat Comms 7, 13052 (2016).

26. Ataman, B. et al. Evolution of Osteocrin as an activity-regulated factor in the primate brain. Nature 539, 242–247 (2016).

27. Luo, G.-Z. et al. Characterization of eukaryotic DNA N6-methyladenine by a highly sensitive restriction enzyme-assisted sequencing. Nat Comms 7, 11301 (2016).

28. Vovis, G. F. & Lacks, S. Complementary action of restriction enzymes endo R · DpnI and endo R · DpnII on bacteriophage f1 DNA. Journal of Molecular Biology 115, 525–538 (1977).

29. Lacks, S. & Greenberg, B. A deoxyribonuclease of Diplococcus pneumoniae specific for methylated DNA. J. Biol. Chem. 250, 4060–4066 (1975).

30. Birnboim, H. C., Sederoff, R. R. & Paterson, M. C. Distribution of Polypyrimidine. Polypurine Segments in DNA from Diverse Organisms. European Journal of Biochemistry 98, 301–307 (1979).

31. Manor, H., Rao, B. S. & Martin, R. G. Abundance and degree of dispersion of genomic d(GA) n ·d(TC) n sequences. Journal of Molecular Evolution 27, 96–101 (1988).

32. Soeller, W. C., Poole, S. J. & Kornberg, T. In vitro transcription of the Drosophila engrailed gene. Genes Dev. 2, 68–81 (1988).

33. Biggin, M. D. & Tjian, R. Transcription factors that activate the Ultrabithorax promoter in developmentally staged extracts. Cell 53, 699–711 (1988).

34. Wallrath, L. L. & Elgin, S. C. Position effect variegation in Drosophila is associated with an altered chromatin structure. Genes Dev. 9, 1263–1277 (1995).

35. Stephens, C., Reisenauer, A., Wright, R. & Shapiro, L. A cell cycle-regulated bacterial DNA methyltransferase is essential for viability. PNAS 93, 1210–1214 (1996).

36. Liu, P. et al. Deficiency in a glutamine-specific methyltransferase for release factor causes mouse embryonic lethality. Mol. Cell. Biol. 30, 4245–4253 (2010).

37. Ghosh, A., Carnahan, J. & Greenberg, M. E. Requirement for BDNF in activity-dependent survival of cortical neurons. Science 263, 1618–1623 (1994).

38. Peters, J., Kalivas, P. W. & Quirk, G. J. Extinction circuits for fear and addiction overlap in prefrontal cortex. Learn. Mem. 16, 279–288 (2009).

39. Lubin, F. D., Roth, T. L. & Sweatt, J. D. Epigenetic regulation of BDNF gene transcription in the consolidation of fear memory. J. Neurosci. 28, 10576–10586 (2008).

40. Aid, T., Kazantseva, A., Piirsoo, M., Palm, K. & Timmusk, T. Mouse and rat BDNF gene structure and expression revisited. J. Neurosci. Res. 85, 525–535 (2007).

41. West, A. E. & Orlando, V. Epigenetics in brain function. Neuroscience 264, 1–3 (2014).

42. West, A. E. Biological functions of activity-dependent transcription revealed. Neuron 60, 523–525 (2008).

43. Sakata, K. et al. Role of activity-dependent BDNF expression in hippocampal-prefrontal cortical regulation of behavioral perseverance. Proc. Natl. Acad. Sci. U.S.A. 110, 15103–15108 (2013).

44. Giresi, P. G., Kim, J., McDaniell, R. M., Iyer, V. R. & Lieb, J. D. FAIRE (Formaldehyde-Assisted Isolation of Regulatory Elements) isolates active regulatory elements from human chromatin. Genome Res. 17, 877–885 (2007).

45. Simon, J. M., Giresi, P. G., Davis, I. J. & Lieb, J. D. Using formaldehyde-assisted isolation of regulatory elements (FAIRE) to isolate active regulatory DNA. Nature Protocols 7, 256–267 (2012).

46. Funahashi, N. et al. YY1 positively regulates human UBIAD1 expression. Biochemical and Biophysical Research Communications 460, 238–244 (2015).

47. Makhlouf, M. et al. A prominent and conserved role for YY1 in Xist transcriptional activation. Nat Comms 5, 4878 (2014).

48. Ratel, D., Ravanat, J.-L., Berger, F. & Wion, D. N6-methyladenine: the other methylated base of DNA. BioEssays 28, 309–315 (2006).

49. Low, D. A., Weyand, N. J. & Mahan, M. J. Roles of DNA adenine methylation in regulating bacterial gene expression and virulence. Infect. Immun. 69, 7197–7204 (2001).

50. Mohammad, F., Mondal, T., Guseva, N., Pandey, G. K. & Kanduri, C. Kcnq1ot1 noncoding RNA mediates transcriptional gene silencing by interacting with Dnmt1. Development 137, 2493–2499 (2010).

51. Di Ruscio, A. et al. DNMT1-interacting RNAs block gene-specific DNA methylation. Nature 503, 371–376 (2013).

52. Wang, Y. et al. WT1 recruits TET2 to regulate its target gene expression and suppress leukemia cell proliferation. Mol. Cell 57, 662–673 (2015).

53. Li, X., Baker-Andresen, D., Zhao, Q., Marshall, V. & Bredy, T. W. Methyl CpG Binding Domain Ultra-Sequencing: a novel method for identifying inter-individual and cell-type-specific variation in DNA methylation. Genes, Brain and Behavior 13, 721–731 (2014).

54. Cheng, X. & Roberts, R. J. AdoMet-dependent methylation, DNA methyltransferases and base flipping. Nucl. Acids Res. 29, 3784–3795 (2001).

55. Diekmann, S. DNA methylation can enhance or induce DNA curvature. The EMBO Journal 6, 4213–4217 (1987).

56. Coulombe, B. & Burton, Z. F. DNA bending and wrapping around RNA polymerase: a ‘revolutionary’ model describing transcriptional mechanisms. Microbiol. Mol. Biol. Rev. 63, 457–478 (1999).

57. Guo, Q., Lu, M. & Kallenbach, N. R. Effect of hemimethylation and methylation of adenine on the structure and stability of model DNA duplexes. Biochemistry (1995).

58. Kellinger, M. W. et al. 5-formylcytosine and 5-carboxylcytosine reduce the rate and substrate specificity of RNA polymerase II transcription. Nat. Struct. Mol. Biol. 19, 831–833 (2012).

59. Wang, L. et al. Molecular basis for 5-carboxycytosine recognition by RNA polymerase II elongation complex. Nature 523, 621–625 (2015).

60. Tian, F. et al. Dynamic chromatin remodeling events in hippocampal neurons are associated with NMDA receptor-mediated activation of Bdnf gene promoter 1. Journal of Neurochemistry 109, 1375–1388 (2009).

## References

1. Jung, M. et al. Longitudinal epigenetic and gene expression profiles analyzed by three-component analysis reveal down-regulation of genes involved in protein translation in human aging. Nucl. Acids Res. 43, e100–e100 (2015).

2. Song, G. & Wang, L. Nuclear Receptor SHP Activates miR-206 Expression via a Cascade Dual Inhibitory Mechanism. PLOS ONE 4, e6880 (2009).

3. Pan, H. et al. Negative Elongation Factor Controls Energy Homeostasis in Cardiomyocytes. Cell Reports 7, 79–85 (2014).

4. Chaudhary, P. et al. HSP70 binding protein 1 (HspBP1) suppresses HIV-1 replication by inhibiting NF-κB mediated activation of viral gene expression. Nucl. Acids Res. 44, 1613–1629 (2016).

5. Luo, G.-Z. et al. Characterization of eukaryotic DNA N6-methyladenine by a highly sensitive restriction enzyme-assisted sequencing. Nat Comms 7, 11301 (2016).

6. Li, H. & Durbin, R. Fast and accurate short read alignment with Burrows-Wheeler transform. Bioinformatics 25, 1754–1760 (2009).

7. Li, H. et al. The Sequence Alignment/Map format and SAMtools. Bioinformatics 25, 2078–2079 (2009).

8. Dunham, I. et al. An integrated encyclopedia of DNA elements in the human genome. Nature 489, 57–74 (2012).

9. Huang, D. W., Sherman, B. T. & Lempicki, R. A. Bioinformatics enrichment tools: paths toward the comprehensive functional analysis of large gene lists. Nucl. Acids Res. 37, 1–13 (2009).

10. Da Wei Huang, Sherman, B. T. & Lempicki, R. A. Systematic and integrative analysis of large gene lists using DAVID bioinformatics resources. Nature Protocols 4, 44–57 (2009).

11. Giresi, P. G., Kim, J., McDaniell, R. M., Iyer, V. R. & Lieb, J. D. FAIRE (Formaldehyde-Assisted Isolation of Regulatory Elements) isolates active regulatory elements from human chromatin. Genome Res. 17, 877–885 (2007).

12. Simon, J. M., Giresi, P. G., Davis, I. J. & Lieb, J. D. Using formaldehyde-assisted isolation of regulatory elements (FAIRE) to isolate active regulatory DNA. Nature Protocols 7, 256–267 (2012).

13. Lin, Q. et al. The brain-specific microRNA miR-128b regulates the formation of fear-extinction memory. Nature Neuroscience 14, 1115–1117 (2011).

14. Lin, Q. et al. The brain-specific microRNA miR-128b regulates the formation of fear-extinction memory. Nature Neuroscience 14, 1115–1117 (2011).

15. Li, X. et al. Neocortical Tet3-mediated accumulation of 5-hydroxymethylcytosine promotes rapid behavioral adaptation. Proc. Natl. Acad. Sci. U.S.A. 111, 7120–7125 (2014).

